# Systematic perturbations of SETD2, NSD1, NSD2, NSD3 and ASH1L reveals their distinct contributions to H3K36 methylation

**DOI:** 10.1101/2023.09.27.559313

**Authors:** Gerry A. Shipman, Reinnier Padilla, Cynthia Horth, Bo Hu, Eric Bareke, Francisca N. Vitorino, Joanna M. Gongora, Benjamin A. Garcia, Chao Lu, Jacek Majewski

**Affiliations:** Department of Human Genetics, McGill University, Montreal, Quebec, H3A 1B1, Canada; McGill University Genome Centre, Montreal, Quebec, H3A 0G1, Canada; Department of Biochemistry & Molecular Biophysics, Washington University School of Medicine, St. Louis, Missouri 63110, USA; Department of Genetics and Development, Columbia University Irving Medical Center, New York, New York 10032, USA

**Keywords:** SETD2, NSD1, NSD2, NSD3, ASH1L, H3K36 methylation, epigenetics, transcription

## Abstract

**Background:** Methylation of histone 3 lysine 36 (H3K36me) has emerged as an essential epigenetic component for the faithful regulation of gene expression. Despite its importance in development, disease, and cancer, how the molecular agents collectively shape the H3K36me landscape is unclear.

**Results:** We use a mouse mesenchymal stem cell model to perturb the H3K36me deposition machinery and infer the activities of the five most prominent players: SETD2, NSD1, NSD2, NSD3, and ASH1L. We find that H3K36me2 is the most abundant of the three methylation states and is predominantly deposited at intergenic regions by NSD1, and partly by NSD2. In contrast, H3K36me1/3 are most abundant within exons and are positively correlated with gene expression. We demonstrate that while SETD2 deposits most H3K36me3, it also deposits H3K36me2 within transcribed genes. Additionally, loss of SETD2 results in an increase of exonic H3K36me1, suggesting other H3K36 methyltransferases (K36MTs) prime gene bodies with lower methylation states ahead of transcription. Through a reductive approach, we uncover the distribution patterns of NSD3- and ASH1L-catalyzed H3K36me2. While NSD1/2 establish broad intergenic H3K36me2 domains, NSD3 deposits H3K36me2 peaks on active promoters and enhancers. Meanwhile, the activity of ASH1L is restricted to the regulatory elements of developmentally relevant genes, and our analyses implicate PBX2 as a potential recruitment factor.

**Conclusions:** Within genes, SETD2 deposits both H3K36me2/3, while the other K36MTs are capable of depositing H3K36me1/2 independently of SETD2 activity. For the deposition of H3K36me1/2, we find a hierarchy of K36MT activities where NSD1>NSD2>NSD3>ASH1L. While NSD1 and NSD2 are responsible for most genome-wide propagation of H3K36me2, the activities of NSD3 and ASH1L are confined to active regulatory elements.

## Background

Epigenetic control of gene expression relies on a fine balance between the deposition and maintenance of a myriad of chromatin modifications. The amino terminal tail of histone 3 is a heavily post-translationally modified region of the nucleosome, with the various modifications having diverse roles and permutations that maintain a balance between heterochromatin, euchromatin, and open chromatin regions. Methylation at the lysine 36 position of histone 3 (H3K36) is known to be associated with transcriptionally active regions of the genome. The highest methylation state, H3K36me3, is a mark of actively transcribed gene bodies (1,2). More recently, the intermediate methylation state, H3K36me2, has garnered increased interest, with evidence supporting its importance in preventing compaction of intergenic regions by restricting expansion of antagonistic silencing marks, such as H3K27me2/3, thereby maintaining a balance between gene expression and silencing (3,4). The lowest methylation state, H3K36me1, has received limited attention (5,6), and its significance remains unclear.

Several constituents governing the deposition of H3K36 methylation (H3K36me) are known. H3K36me3, which exists almost entirely within transcribed gene bodies, is deposited by transcriptional coupling of the methyltransferase SETD2 to the elongating RNA polymerase II complex (RNAPII) (7,8). In simple eukaryotes, such as yeast, the presence of H3K36me3 itself has been shown to be important for repressing cryptic transcription, facilitating histone turnover, and regulating pre-mRNA splicing (9,10,11). In contrast, H3K36 mono- and di-methylation are the products of at least four other histone methyltransferases, NSD1/2/3, and possibly ASH1L. While all four of these enzymes have demonstrated H3K36 methyltransferase activity, they appear to have non-redundant functional properties, and their implications in genetic disease and cancer are markedly different. NSD1 loss of function mutations have been identified in most patients with Sotos syndrome, a developmental overgrowth disorder, and also in a subgroup of HPV-negative head and neck squamous cell carcinomas (HNSCC). Here, loss of NSD1 results in depletion of intergenic H3K36me2, and a concomitant decrease of H3K27ac and DNA methylation (DNAme), thereby reducing the activity of associated cis-regulatory elements (CREs) and the expression of their putative target genes (12,13,14,15). In contrast, NSD2 truncating and missense variants have been causally associated with Wolf-Hirschhorn syndrome, a genetic disorder characterized by intellectual and developmental delay (16,17,18). Translocation-mediated overexpression and hyperactive variants of NSD2 and NSD3 have been identified in several types of cancer, notably in multiple myeloma, lung squamous cell carcinoma, and pancreatic ductal adenocarcinoma, with elevated H3K36me2 being implicated in tumour progression and survival outcomes (19,20,21). As an H3K36 methyltransferase, ASH1L remains relatively understudied. However, nonsense and missense variants have been linked with autism spectrum disorder and brain development, and its overexpression has been identified in both anaplastic thyroid cancer and acute leukemia (22,23,24).

H3K36me is an essential epigenetic modification for the establishment and maintenance of gene regulatory programs, and impacts the deposition of many other histone marks, as well as DNAme (25,26,27,28). Clearly, there are considerable pathological consequences when it is aberrantly regulated, underscoring the necessity to better understand its functional significance. Here, we use a mouse mesenchymal stem cell model to dissect the roles of the most well-established H3K36 methyltransferases (K36MTs) in the deposition and maintenance of this modification. Mesenchymal stem cells, which are multipotent, can be induced into oncogenic transformation (29,30,31). In this study, we use CRISPR-Cas9 gene editing to assemble a panel of single and combinatorial knockout cell lines, allowing us to deconstruct the individual contributions of each methyltransferase to the overall H3K36me landscape. Our work provides new insights into the distinct and overlapping genomic preferences of lysine 36 methyltransferases and consolidates existing knowledge in the context of a structured analysis.

## Results

### H3K36 methylation states have distinct distribution patterns

In order to characterize the effects of individual K36MTs in shaping the H3K36me landscape, we first sought to describe the deposition patterns of H3K36me in wild-type C3H10T1/2 mouse mesenchymal stem cells (mMSCs). Therefore, we used liquid chromatography - tandem mass spectrometry (MS) to quantify the genome-wide abundance and chromatin immunoprecipitation with sequencing (ChIP-seq) to profile the genomic distribution of all three H3K36me states. Consistent with previous observations in this cell type, we find that H3K36me2 is broadly distributed both within genes and intergenic regions (IGRs), while H3K36me3 is restricted specifically to actively transcribed genes (Fig. 1a) (4,11). Interestingly, we find that H3K36me1 is also found predominantly within genes, with lower levels also found in IGRs (Fig. 1a). As quantified by MS, H3K36me2 is the most abundant of the three methylation states, marking approximately 30% of all H3 peptides (Fig. 1b). In comparison, H3K36me1 and H3K36me3 occupy approximately 14% and 7% of H3 peptides, respectively (Fig. 1b).

**Fig. 1.**
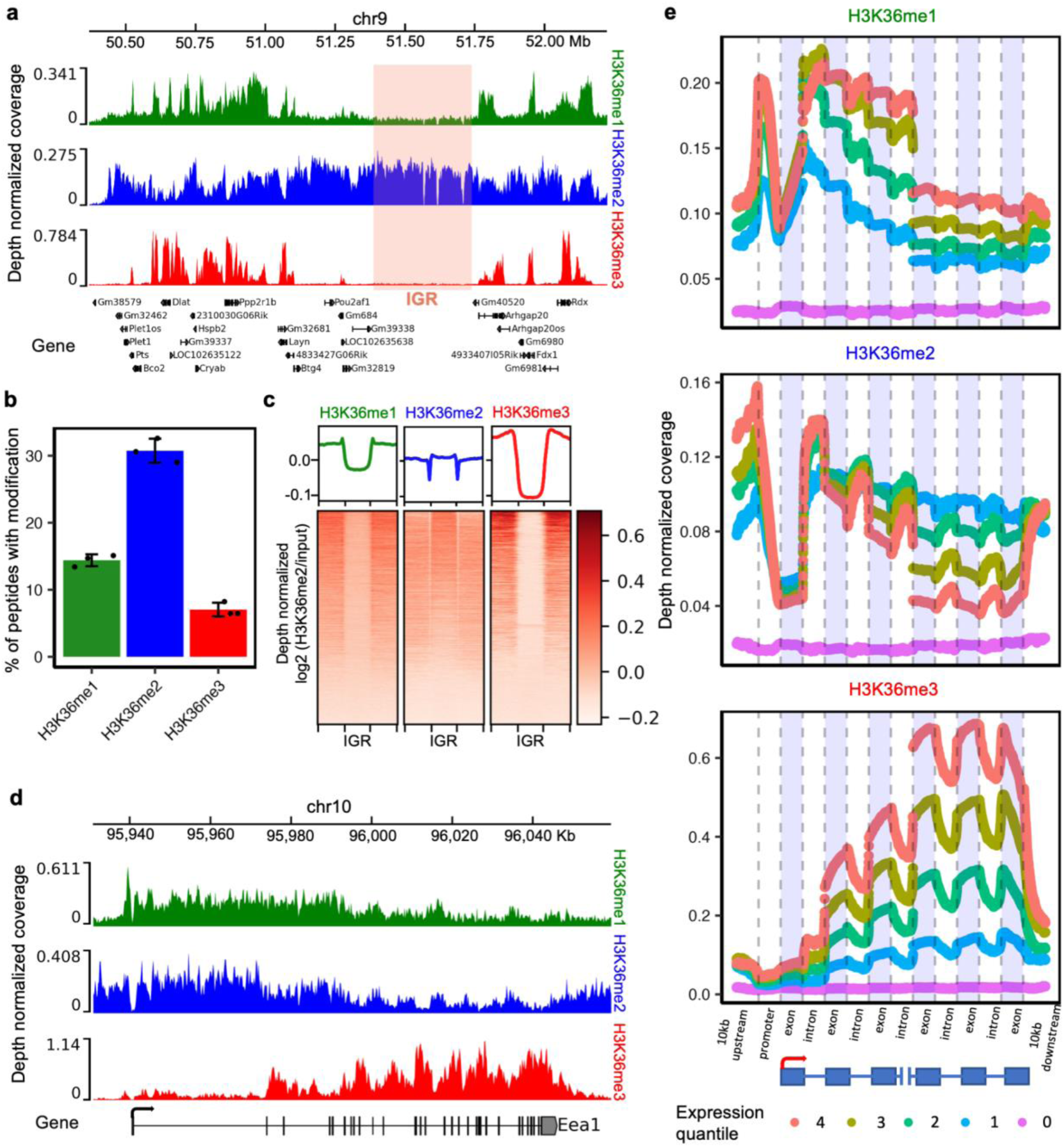
The three H3K36me states have distinct genome-wide distributions. **a.** Genome-browser representation of the three H3K36me states in wildtype C3H10T1/2 mMSCs, illustrating the broad distribution of H3K36me2 in genic and intergenic regions, while H3K36me1 and H3K36me3 are predominantly distributed within genic regions. **b.** Bulk levels of H3K36me1/2/3 as quantified by MS, with H3K36me2 showing the highest abundance followed by H3K36me1 and H3K36me3. The error bars indicate mean ± standard deviation. Individual data points represent biological replicates (n=3 per methylation state). **c.** Heatmaps showing H3K36me2 (input-and depth-normalized) enrichment patterns ± 20 kb flanking intergenic regions (IGRs), indicating that at IGRs (n = 9551), there are high levels of H3K36me2, low levels of H3K36me1 and lack of H3K36me3. **d.** Genome-browser coverage tracks demonstrating that H3K36me1/2 are highest at 5’ regions, and then decrease towards the 3’ end, where they are replaced by H3K36me3 in regions of high exon density. **e.** Representative view of genic H3K36me1/2/3 distributions, highlighting the 5’ to 3’ distribution patterns, exonic versus intronic signal, and dependence on gene expression for the three H3K36 methylation states. Only transcripts with at least 50000 base-pairs (bp) and 6 exons were used in the analysis. An aggregate of H3K36me signals on the first three exons and the last three exons are shown. The expression quantiles were calculated based on normalized (reads per kilobases) expression counts from the parental samples. Expression quantile 4 comprises transcripts with the highest expression, expression quantile 1 comprises transcripts with the lowest expression greater than zero, and expression quantile 0 comprises transcripts with zero counts.

To assess the global distribution and abundance of H3K36me within IGRs, we centered ChIP-seq signals on these regions to generate aggregate profiles. Expectedly, we find that H3K36me3 is essentially absent and confined to flanking genic regions (Fig. 1c). In contrast, the levels of H3K36me2 are greater within IGRs, reflecting its functional importance within these regions (Fig. 1c) (4), while low levels of H3K36me1 are also found within IGRs.

Upon examination of genome browser coverage tracks within transcribed genes, we find that the levels of H3K36me1 and H3K36me2 are highest in 5’ regions, plateau within the first intron, and then decrease towards the 3’ end, where they are replaced by H3K36me3 (Fig. 1d). To closely examine the gene body distribution patterns of H3K36me, we divided actively transcribed genes into expression quantiles and assessed the differences of individual H3K36me marks within introns and exons across varying levels of gene expression (Fig. 1e). We find that bulk enrichment of H3K36me1 and H3K36me3 signals are greater in exons compared to introns, whereas the inverse relationship is true for H3K36me2 (Fig. S1a). Although the 5’ to 3’ profiles of H3K36me1 and H3K36me2 are similar, they correlate differently with gene expression levels: H3K36me1 and H3K36me3 are positively correlated with gene expression whereas H3K36me2 is not, likely reflecting the conversion efficiency of H3K36me2 to H3K36me3 within the exons of highly transcribed genes (Fig. S1b).

In summary, the dynamics of the three methylation states within gene bodies are influenced by the levels of transcription: H3K36me1/2 decrease from 5’ to 3’ in gene bodies where they are replaced by H3K36me3; H3K36me1/3 are positively correlated with gene expression, whereas H3K36me2 is not; and H3K36me2 is the most abundant methylation state and is broadly distributed in IGRs, whereas H3K36me1/3 are predominantly found within genes.

### Individual and combined H3K36 methyltransferase contributions to bulk H3K36me

Each of the most well-established K36MTs - SETD2, NSD1, NSD2, NSD3 and ASH1L - have been shown biochemically to exhibit methyltransferase activity specific for H3K36 via their catalytic Su(var)3-9, Enhancer-of-zeste and Trithorax (SET) domains (9,32,33,34). However, their distinct contributions to the H3K36me landscape is currently unknown. To better understand and decompose these contributions, we used CRISPR- Cas9 gene editing to establish a panel of single and combinatorial knockout mMSC lines, targeting the SET domain of each K36MT (Fig. S2). We used MS and Western blots to quantify the overall changes to bulk levels of H3K36me across different conditions. As described, H3K36me3 is the least abundant methylation state in parental cells, marking approximately 7% of all H3 peptides (Fig. 1a), and consistent with previous reports in mMSCs, it is almost entirely deposited by SETD2 (31). As shown by MS, cells lacking functional SETD2 (SETD2-KO) have at least a 6 fold-reduction of H3K36me3 in comparison to parental cells (Fig. 2a, Fig. S3a). Interestingly, in SETD2-KO cells, the levels of H3K36me1/2 increase relative to parental cells, which is consistent with fewer H3K36 residues being upgraded to higher orders of methylation (Fig. 2a).

**Fig. 2.**
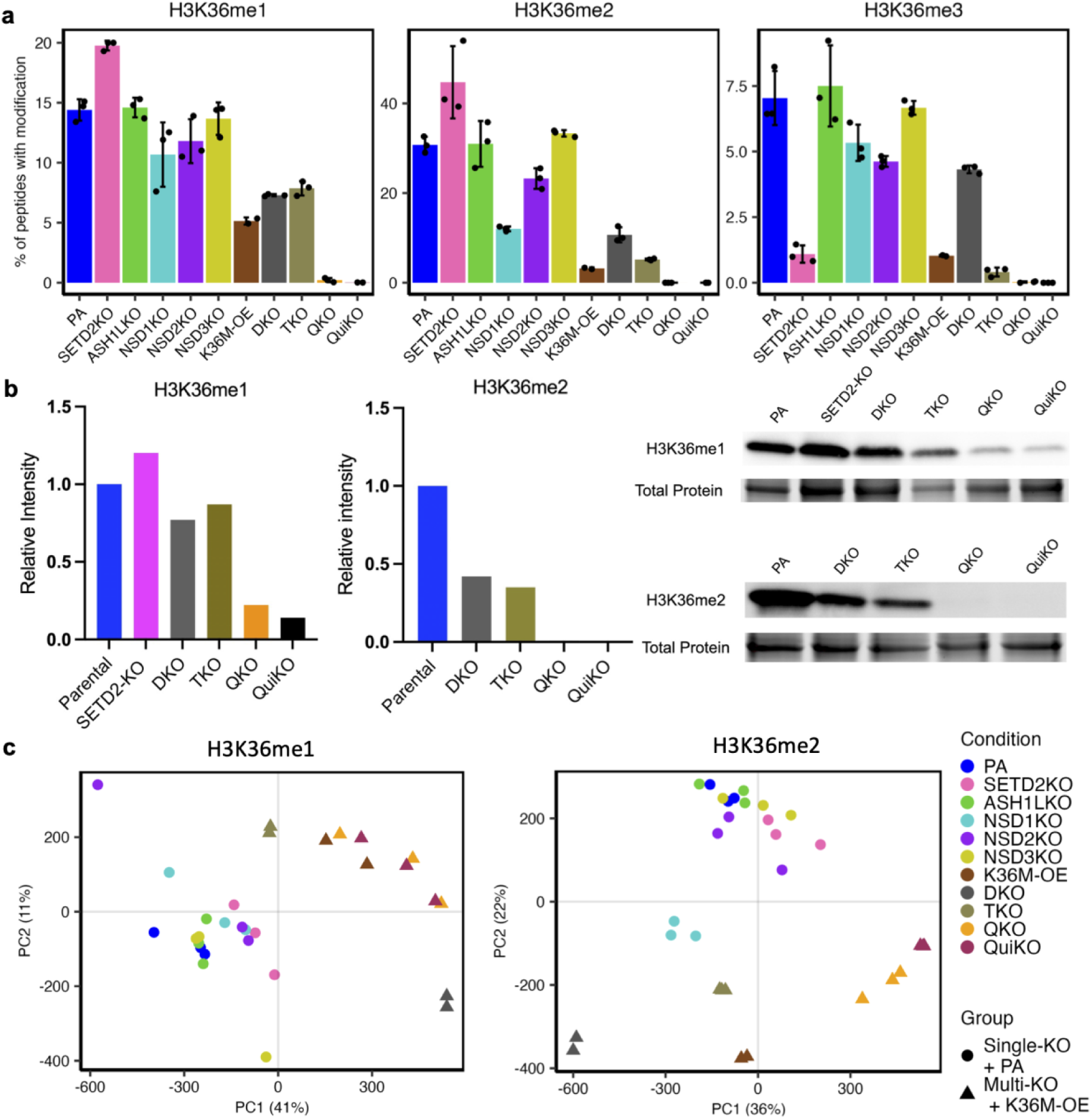
Distinct alterations in H3K36me global abundance are found following single and combinatorial knockouts of K36MTs. **a.** Genome-wide changes to H3K36me1/2/3 in various conditions as quantified by MS. Individual dots denote biological replicates within a condition (n=3 per condition, except for QuiKO where an outlier was omitted for H3K36me1). Error bars indicate mean ± standard deviation. **b.** Quantitative Western blots demonstrating changes to H3K36me1 (left) and H3K36me2 (right) in the various knockout conditions relative to parental cells. Immunoblot and protein gel cropped for relevant regions. Full total protein loading gel used for quantification of relative intensity illustrated in Fig.S3b. **c.** Principal component analysis (PCA) plot of depth-normalized log2 H3K36me ChIP-Seq signal over input based on 10 kilobase (kb) binned signals (after scaling and centering). Bins with raw read counts consistently lower than 100 across all samples were excluded. Each data point denotes a biological replicate. The colors represent the different conditions. Circles represent the parental and single-KO samples, and the triangles represent the multiple-KO samples and H3K36M-OE.

To investigate the individual contributions of each K36MT to global levels of H3K36me1/2, we generated individual NSD1, NSD2, NSD3, and ASH1L knockout cells, with three biological replicates for each cell line (Fig. S2a). Strikingly, ablation of NSD3 and ASH1L appears to have no discernible effect on the global abundance of H3K36me1/2 (Fig. 2a). While both of these enzymes have previously been reported to deposit these modifications (33,34), these results suggest that their effects may be restricted to specific loci or masked by the effects of other K36MTs, such as NSD1 or NSD2. In contrast, loss of NSD1 results in the greatest depletion of both H3K36me1/2, especially H3K36me2 which is reduced more than 2-fold, while in NSD2-KO cells, H3K36me2 levels are reduced by approximately 12% (Fig. 2a).

Further decreases of H3K36me2 are observed in multi-knockout conditions. The concurrent loss of NSD1 and NSD2 (DKO) in mMSCs leads to a 3-fold reduction in global H3K36me2 levels (Fig. 2a/b, Fig. S3a), which is consistent with previous studies (31). In triple knockout cells lacking NSD1/2-SETD2 (hereafter referred to as TKO), bulk levels of H3K36me2 are further reduced, indicating that SETD2 also contributes to the deposition of this mark (Fig. 2a/b, Fig. S3a). Similar to SETD2-KO cells, TKO cells are nearly devoid of H3K36me3. Ablation of NSD3 in NSD1/2/3-SETD2 quadruple knockout cells and subsequently ASH1L in NSD1/2/3-SETD2-ASH1L quintuple knockout cells (hereafter referred to as QKO and QuiKO, respectively) results in a total loss of H3K36me2 (Fig. 2a/b), and these observations are further supported by ChIP-Rx (Fig. S3a).

We have previously explored and characterized mMSCs overexpressing the H3K36M oncohistone (H3K36M-OE), a mutation frequently identified in chondroblastoma patients, and found that one of the primary effects of this mutation is global H3K36 hypomethylation (31). Comparing the effects of this mutation to our combinatorial knockouts, we find that the lowest levels of H3K36me are observed in the H3K36M-OE, QKO, and QuiKO cell lines, confirming that this mutation exerts a global effect on the activity of multiple K36MTs (Fig. 2a, Fig. S3a). Indeed, principal component analysis (PCA) of the three methylation states reveals that cells with multiple K36MT deletions cluster more closely with cells overexpressing the H3K36M mutation (Fig. 2c, Fig. S3c). Interestingly, for H3K36me2, the NSD1-KO samples also cluster more closely with the multi-knockout and H3K36M-OE samples, supporting that NSD1 deposits the bulk of H3K36me2. In comparison, the other individual knockout conditions cluster more closely with parental cells (Fig. 2c, Fig. S3c).

Overall, as it pertains to the global abundance of the three H3K36me states in mMSCs, SETD2 deposits nearly all H3K36me3, while NSD1 is responsible for the majority of bulk H3K36me1/2 levels. In comparison, NSD2 also deposits modest amounts of H3K36me1/2, while NSD3 and ASH1L have the smallest contributions.

### H3K36me1 is primarily associated with gene bodies

In both disease and development, nearly all previous investigations of H3K36me and its respective writers have focused on the higher orders of methylation, H3K36me2/3, with H3K36me1 remaining comparatively understudied. As revealed by MS and ChIP-seq, the global abundance of H3K36me1 is significantly reduced in both NSD1/2-DKO and H3K36M-OE cells (Fig. 2a, Fig. 3a). In QKO and QuiKO cells, H3K36me1 is virtually absent. However, close analysis of the ChIP-seq profiles demonstrates that the distribution pattern of the extremely low levels of remaining H3K36me1 closely resembles that of parental cells (Fig. 2b, Fig. 3a). Given that the five most well-established K36MTs are ablated in the QuiKO condition, this intriguingly suggests that there may be another K36MT capable of depositing this modification.

**Fig. 3.**
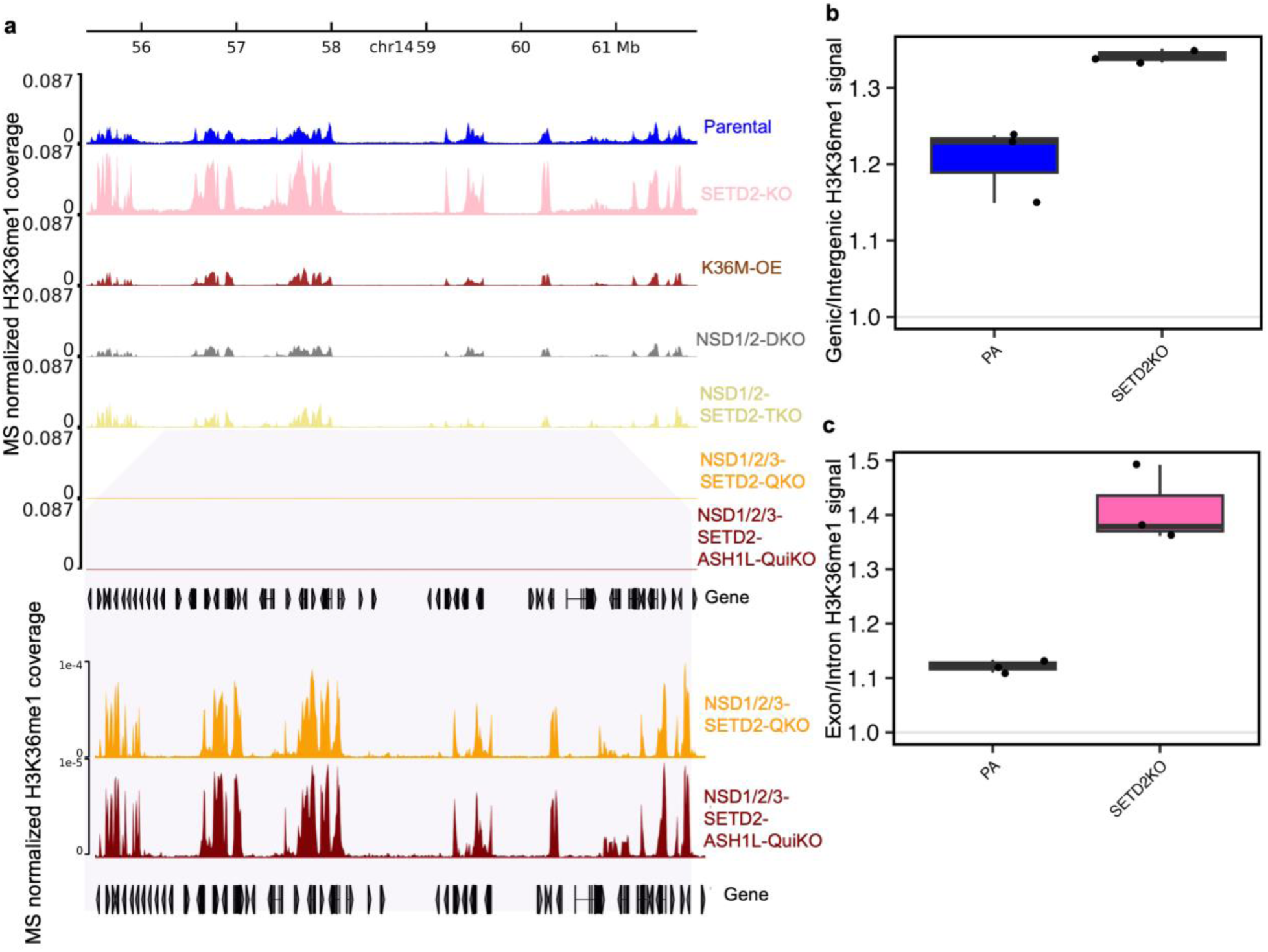
Effects of K36MT ablations and H3K36M-OE on H3K36me1 distribution patterns within genes. **a** Genome-browser tracks of MS-normalized H3K36me1 signal showing depletion in H3K36M-OE and multi-knockout conditions, and an increase in SETD2-KO cells. The two bottom tracks illustrates that despite NSD1/2/3-SETD2-QKO and NSD1/2/3-SETD2-ASH1L-QuiKO displaying reduced amounts of H3K36me1, the distribution patterns remain similar to that observed in parental cells. **b**. Boxplots of H3K36me1 genic to intergenic ratios, indicating that the increase in SETD2-KO cells is found primarily within genic regions. (n=3 per condition) **c**. Boxplots showing the exon versus intron H3K36me1 signal in parental and SETD2-KO cells, indicating increased signal at exons following SETD2-KO. (n=3 per condition).

Interestingly, in SETD2-KO cells, the global abundance of H3K36me1 increases (Fig. 2a/b, Fig. 3a). We hypothesized that the increase of H3K36me1 after loss of SETD2 is, at least in part, due to mono-methylated peptides no longer being upgraded to higher orders of methylation. Since SETD2 is known to act primarily within gene bodies, we assessed whether the increase of H3K36me1 occurs primarily within genes or IGRs. Indeed, in comparing the genic to intergenic ratios of H3K36me1 signal between parental and SETD2-KO cells, we find that the increase of H3K36me1 occurs predominantly within genes (Fig. 3b). Subsequent in-depth analysis of the changes within gene bodies reveals that the increase of H3K36me1 occurs primarily within exons in comparison to introns (Fig. 3c). Overall, these results support that when the activity of SETD2 is lost, the cascade of methylation from H3K36me1 to H3K36me3 is disrupted primarily in genes, and more specifically, within exons, leading to an increase of H3K36me1 in these regions. Furthermore, these results suggest that other K36MTs may prime gene bodies with H3K36me1 in advance of SETD2 activity, which is tightly associated with transcription.

### Intergenic H3K36me2 is predominantly deposited by NSD1 and in part by NSD2 in mMSCs

We and others have previously found that one of the predominant effects of the H3K36M mutation is a marked reduction of intergenic H3K36me2, an effect that can be largely recapitulated from the combined loss of NSD1 and NSD2 (Fig. 4a) (31). In this context, however, the individual contributions of NSD1 and NSD2 to the intergenic H3K36me2 landscape in mMSCs remains unclear. Therefore, we evaluated changes to intergenic H3K36me2 in both NSD1-KO and NSD2-KO cells. Consistent with reports in other cell types (4,15), loss of NSD1 results in the greatest depletion of intergenic H3K36me2, reducing it nearly 2-fold (Fig. 4a, Fig. 4b). In contrast, the loss of NSD2 also reduces intergenic H3K36me2, although to a much lower extent. Comparing these results to NSD1/2-DKO cells, we report that while both NSD1-KO and NSD2-KO cells have an effect on intergenic H3K36me2, neither condition alone can recapitulate the depletion observed in DKO cells, suggesting that NSD1 and NSD2 may propagate this mark in IGRs through an additive effect, with NSD1 playing the predominant role (Fig. 4a, Fig. 4b). In comparing the effects of the other K36MTs, no discernible changes to intergenic H3K36me2 are observed in SETD2-KO, NSD3-KO, or ASH1L-KO cells (Fig.S4a).

**Fig. 4.**
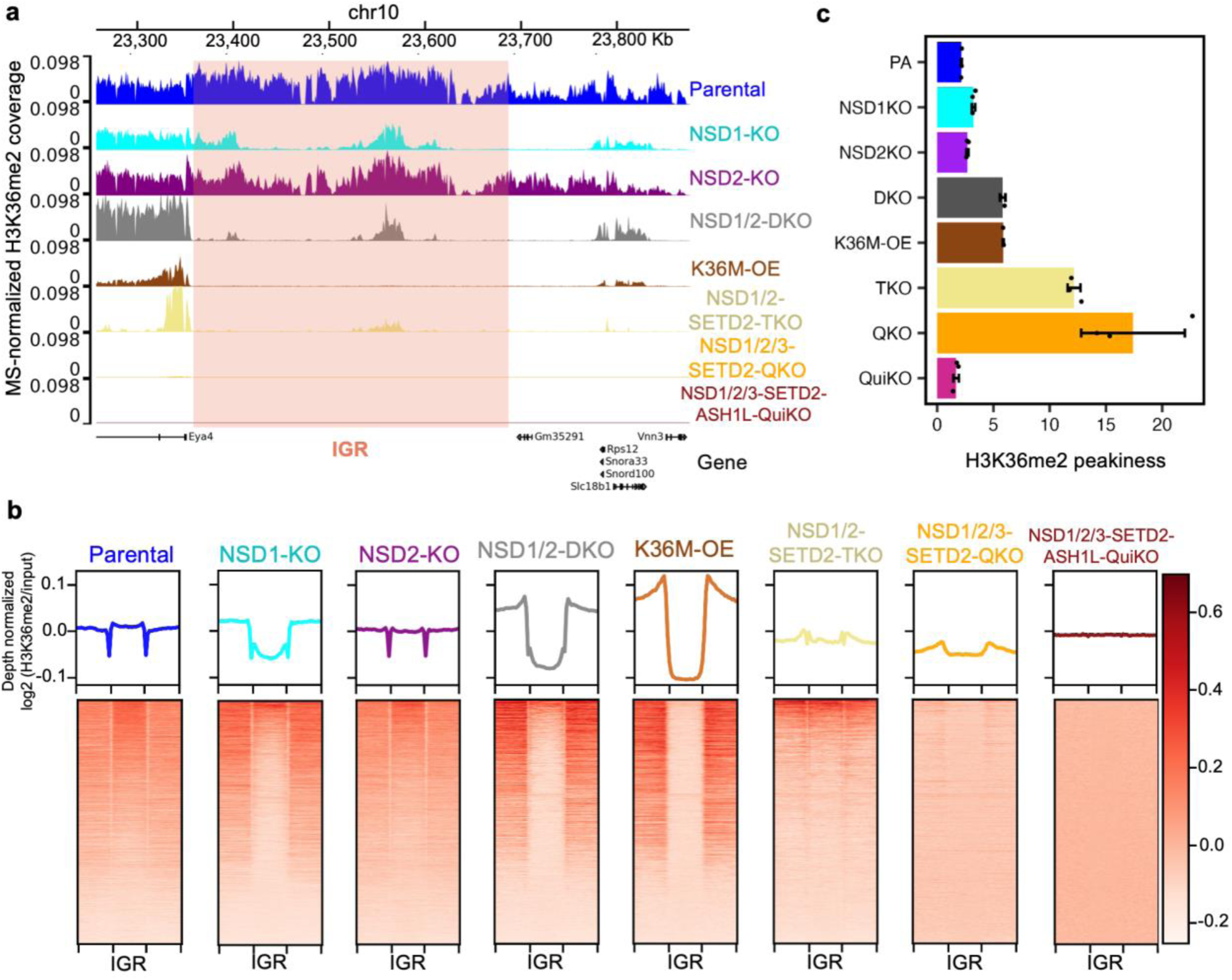
Intergenic H3K36me2 decreases following NSD1-KO, and is further reduced in H3K36M-OE and multi-knockout conditions. **a.** Coverage tracks of MS-normalized H3K36me2 signal centered on intergenic regions flanked by genic regions, highlighting reduced intergenic signal following loss of NSD1, more pronounced reductions in NSD1/2-DKO and H3K36M-OE cells, and the most significant depletions in subsequent multi-knockout conditions. **b.** Heatmaps showing H3K36me2 (input- and depth-normalized) enrichment patterns ± 20 kb flanking IGRs, summarizing the H3K36me2 trends for multiple cell lines at IGRs. **c**. H3K36me2 “peakiness scores”, representing the average ChIP signal in the top 1% of 1-kb bins over the total signal, indicating more punctate distributions of H3K36me2 in multi-knockout conditions. Individual data points represent biological replicates for each condition (n=3 per condition, except for DKO and K36M-OE where n=2). Error bars represent mean ± standard deviation.

Intergenic H3K36me2 domains span megabases and are essential components of the gene regulatory network, often demarcating enhancers and other CREs (15). Therefore, it is worth noting that while most of the individual K36MTs do not exert a global effect on the broad intergenic distribution of H3K36me2, the distribution does become confined to smaller regions upon loss of NSD1 (Fig.4a, Fig.4c), supporting that NSD1 is the predominant K36MT in these regions and that the other K36MTs may act with more specificity. Indeed, even in the absence of NSD1, the combined catalytic activities of the other K36MTs are insufficient to rescue the loss of these broad H3K36me2 domains (Fig.4a, Fig.4b). The H3K36me2 signal is further attenuated in DKO cells, and becomes progressively more punctate in subsequent multi-knockout conditions (i.e. in TKO, QKO, and QuiKO cells). To visualize this, we generated “peakiness” scores for all KO conditions, which represent the average H3K36me2 ChIP signal in the top 1% of 1-kb bins and where a higher score indicates a more punctate distribution (Fig.4c, Fig.S4b).

Taken together, NSD1 is consistently the primary K36MT responsible for the establishment and maintenance of broad intergenic H3K36me2 domains, while in mMSCs where it is expressed less, NSD2 also significantly contributes to the deposition of this mark.

### Genic H3K36me distribution patterns are highly dependent on SETD2 occupancy

In the presence of the H3K36M mutation, which affects all K36MTs including SETD2, the profile of H3K36me2 within genes exhibits a shift towards 3’ regions and becomes elevated within exons, as compared to introns (Fig. 5a-d). In NSD1/2-DKO cells, we found a similar trend, where although the preference for the 5’ region of genes remains unchanged, the absence of NSD1/2 results in a remarkable change in the dependence on expression levels: H3K36me2 is now positively correlated with gene activity (Fig. 5b- c). While the distribution of H3K36me2 in the highly transcribed genes of DKO cells remains higher within introns than exons, in lowly transcribed genes there is now greater signal in exons compared to introns, similar to the distribution pattern of H3K36me3 (Fig. 1e), which may result from reduced occupancy of SETD2 in genes transcribed at lower frequencies (Fig. 5b). The observed distinction between the highly and lowly transcribed genes in DKO cells can likely be attributed to the increased prominence of SETD2 as the primary K36MT for H3K36me2 in genic regions, leading to the conversion of H3K36me2 to H3K36me3, specifically in highly transcribed genes. In genes transcribed with less frequency, however, SETD2 may primarily deposit H3K36me2 without further conversion. Furthermore, in SETD2-KO cells, H3K36me2 is no longer converted to H3K36me3, and a slight positive shift in the correlation between H3K36me2 and gene expression is observed (Fig. 5b, Fig. 5c). In contrast, the individual knockouts of ASH1L, NSD1, NSD2 and NSD3 do not change the dependency of H3K36me2 on gene expression (Fig. S4c). Nonetheless, the strong positive correlations between H3K36me2 and gene expression quantiles observed in H3K36M-OE, NSD1/2-DKO, and SETD2-KO cells are consistently observed when assessing genes on a genome-wide scale without segregating them based on quantiles, and in this context the overall signal demonstrates an exonic enrichment in comparison to introns. (Fig. 5d, Fig. S4d).

**Fig. 5.**
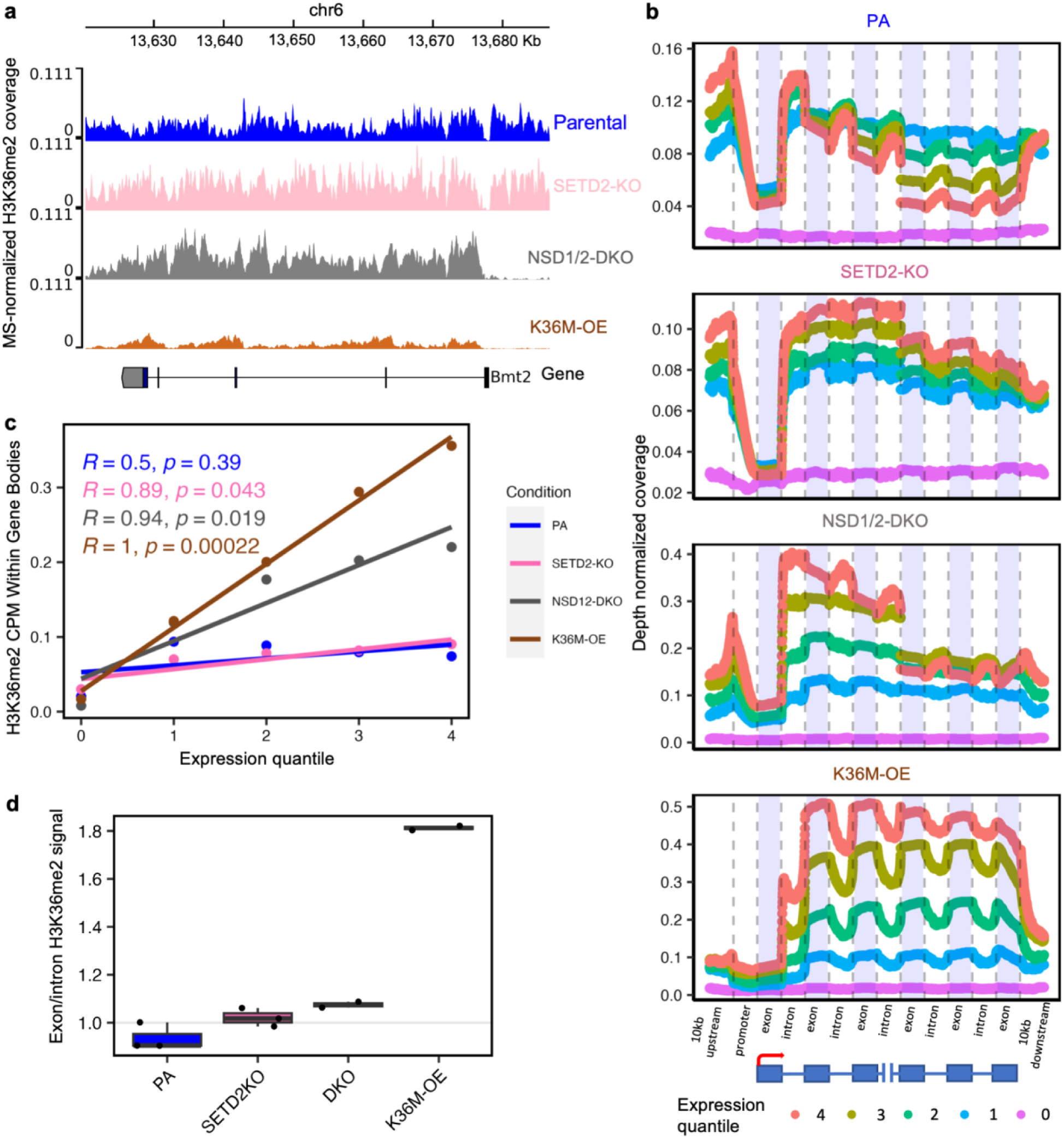
Genic H3K36me2 distribution patterns are altered following SETD2-KO, NSD1/2- DKO, and H3K36M-OE. **a.** Coverage tracks of MS-normalized H3K36me2 signal showing changes in genic distribution following specific K36MT knockouts. **b**. Zoomed in view of H3K36me2 distribution in gene bodies, highlighting changes in exon versus intron signal and dependence on gene expression following K36MT deletions. Only transcripts with at least 50000 base-pairs (bp) and 6 exons were used in the analysis. An aggregate of H3K36me signals on the first three exons and the last three exons are shown. The expression quantiles were calculated based on normalized (reads per kilobases) expression counts from the parental samples. Expression quantile 4 comprises transcripts with the highest expression, expression quantile 1 comprises transcripts with the lowest expression greater than zero whereas expression quantile 0 comprises transcripts with zero counts. **c.** Correlation plots of H3K36me2 CPM and gene expression quantiles, indicating a shift to significant positive correlations between H3K36me2 and gene expression following SETD2-KO, NSD12-DKO and H3K36M-OE. **d.** Boxplots of mean H3K36me2 signal at exons versus introns, indicating a trend towards greater signal at exons compared to introns following SETD2-KO, NSD12-DKO and H3K36M-OE.

### NSD3-mediated H3K36me2 is targeted to active regulatory regions

Although depletion of NSD1, NSD2 and SETD2 results in a progressive loss of H3K36me2, with NSD1/2 predominantly affecting intergenic regions, and SETD2 affecting gene-bodies, a non-negligible amount of H3K36me2 still remains in TKO cells (Fig. 2a, Fig. 6a). The profile of H3K36me2 becomes progressively more punctate, starting from megabase size euchromatic domains in parental cells, and ending with kilobase size broad “peaks” in the TKO cells (Fig. 6a). Close examination of the distribution in TKO cells reveals that most of these residual peaks are centered on TSS and enhancer regions (Fig. 6b, Fig. 6c). Adding more epigenetic information, we note that the majority of these broad peaks straddle regions of open chromatin, as demonstrated by ATAC-seq and the presence of H3K27ac (Fig. 6b, Fig. 6c).

**Fig. 6.**
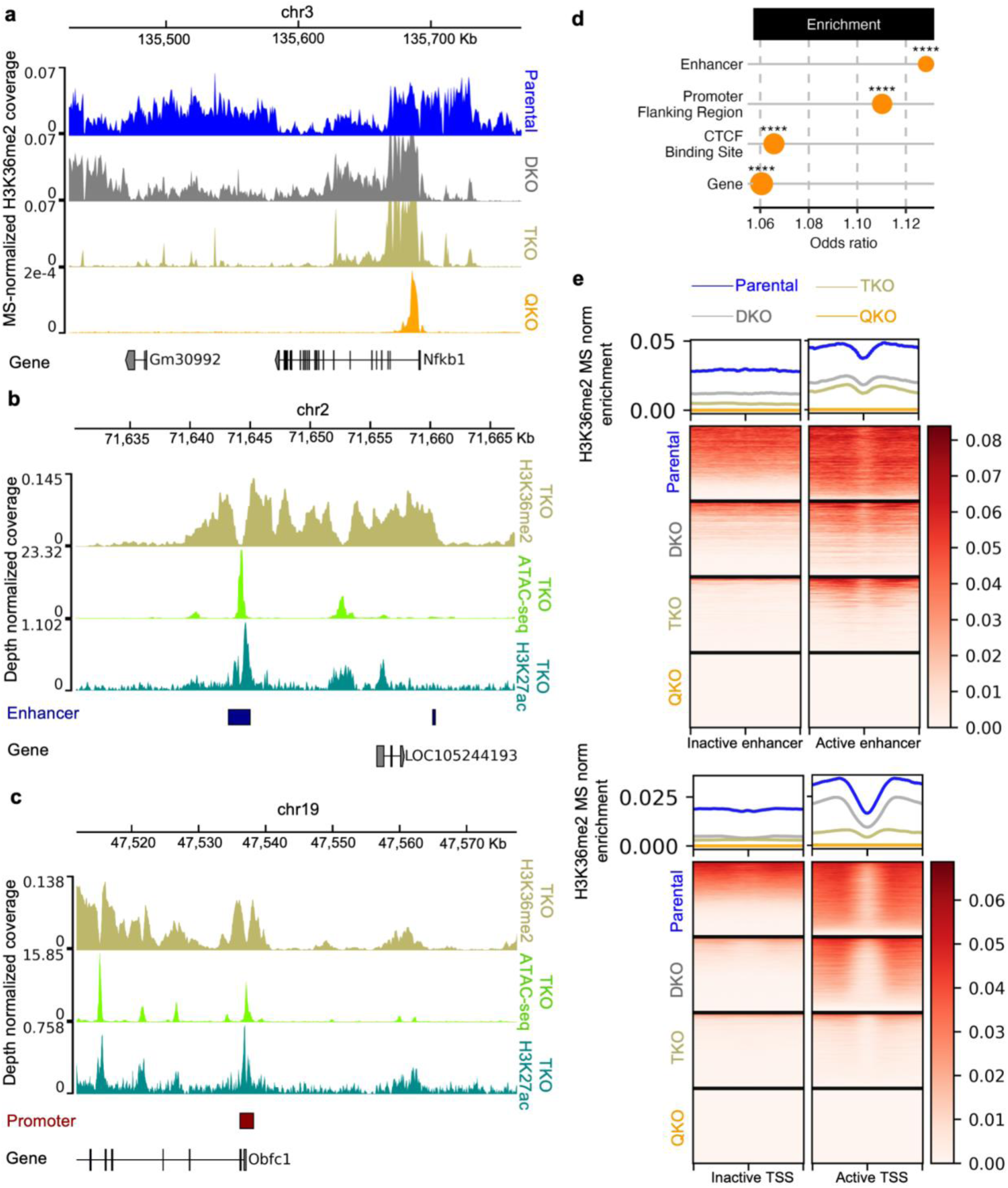
NSD3 deposits H3K36me2 specifically at active promoters and enhancers. **a.** Genome-browser tracks displaying MS-normalized H3K36me2 signal where the distribution becomes more punctate with additional K36MT deletions. **b.** Genome-browser coverage tracks showing NSD1/2-SETD2-TKO (TKO) H3K36me2 signal at active enhancer regions, which are identified by the presence of ATAC-seq and H3K27ac peaks. **c.** Genome-browser coverage tracks displaying H3K36me2 signal for NSD1/2- SETD2-TKO (TKO) at active promoter regions, which are identified by the presence of ATAC-seq and H3K27ac peaks. **d.** Overlap enrichment results of Ensembl annotations with bins found in TKO but not in QKO (i.e., identified as NSD3-catalyzed H3K36me2). The size of the dots corresponds to the number of bins overlapping the corresponding annotation. **** represents p-value < 1e-4 whereas *** represents p-value < 1e-3 based on Fisher’s exact test of bins overlapping a specific class of annotated regions versus a background of all bins in TKO. Only the top four most significant annotations are shown. **e.** Heatmaps showing H3K36me2 (input- and depth-normalized as well as MS-scaled) signal centered on transcription start site (TSS) in inactive (n=18,290) and active promoters (n=18,290) as well as H3K36me2 signal centered on inactive (n=1758) and active enhancers (n=1758), highlighting the loss of H3K36me2 signal at these active regions following NSD3 knockout.

In a wildtype context, it is currently unknown where NSD3 exerts its catalytic activity and to what extent. As previously described, in NSD3-KO cells there is almost no discernible change to the distribution or bulk levels of H3K36me2 (Fig. 2a & Fig. S4a). Therefore, we hypothesized that the effects of NSD3 may be concealed by the activity of other K36MTs. In QKO cells, lacking NSD1/2/3-SETD2, nearly all remaining H3K36me2 is depleted (Fig. 6a), indicating that the residual peaks remaining in TKO cells are deposited by NSD3. We used genomic element annotations and carried out enrichment analysis to characterize the NSD3-deposited H3K36me2 regions (35). Confirming our previous observations, we find that the strongest functional H3K36me2 enrichment categories are at enhancer and promoter-flanking regions (Fig. 6d). After centering MS-normalized H3K36me2 signal on annotated enhancers and TSS’, we find that the signal is abolished following loss of NSD3 in QKO cells (Fig. 6e). Thus, it appears that the residual H3K36me2 found in TKO cells is deposited by NSD3 and its activity is targeted specifically to active promoters and enhancers.

This finding implicates NSD3 in the targeted deposition of H3K36me2 around active regulatory elements (Fig. 6b-e). Because of the broad distribution of H3K36me2 in parental cells, this role only becomes evident in the absence of more catalytically active K36MTs. Indeed, in NSD3-KO cells, it appears that a compensatory effect may exist, where one or more of the other K36MTs may act to deposit H3K36me2 in regions where NSD3 would generally localize. Thus, while NSD3 is able to deposit substantial levels of H3K36me2, its mode of activity differs from NSD1/2: NSD1/2 establish broad H3K36me2 domains both within genes and IGRs, while NSD3 establishes more localized broad peaks centered on promoters and enhancers.

### ASH1L selectively deposits H3K36me2 at the regulatory elements of developmentally relevant genes

Previous studies of ASH1L have demonstrated that it is capable of depositing H3K36me2, and that it has been found to be enriched at the promoters of certain developmentally important genes (34,36). However, its global contributions to H3K36me2 and the precise genomic regions at which it exerts its catalytic activity remains to be elucidated. As described, we depleted ASH1L in parental mMSCs, and detected no discernible changes to the bulk levels or distribution of H3K36me2 (Fig. 2a and Fig. S4a). The QKO cells, lacking NSD1/2/3 and SETD2, exhibit extremely low levels of H3K36me2, reduced by nearly 30x compared to parental cells (Fig. 2a), however, we detected a few (n=119) clearly defined broad (10-70 kb) peaks remaining across the genome (Fig. 7a). These peaks are again centered on regions of open chromatin accessibility, as determined by ATAC-seq, with approximately 60 promoter and 20 deep intergenic sites (Fig. 7a, Fig. 7b). Based on our previous findings, we hypothesized that these remaining peaks are deposited by ASH1L, and indeed, depletion of ASH1L in the QuiKO cells results in a total loss of all remaining H3K36me2 (Fig. 2a, Fig. 7a). It is remarkable that ASH1L exhibits such high specificity for a few selected regulatory regions. To identify possible associations between ASH1L and known DNA binding factors, we carried out motif analysis under the ATAC-seq peaks within the ASH1L-associated regions. We identified enrichment of binding motifs for several developmentally important transcription factors (Fig. 7c). Using mMSC RNA-seq data (37), we find that PBX2 is the only one of these transcription factors that is expressed, implicating PBX2 as the best candidate for recruiting ASH1L to these regions (Fig. 7c, Fig. S5a). Overall, in mMSCs, ASH1L appears to be the least prolific of the K36MTs, and exhibits high specificity for its recruitment to regulatory elements.

**Fig. 7.**
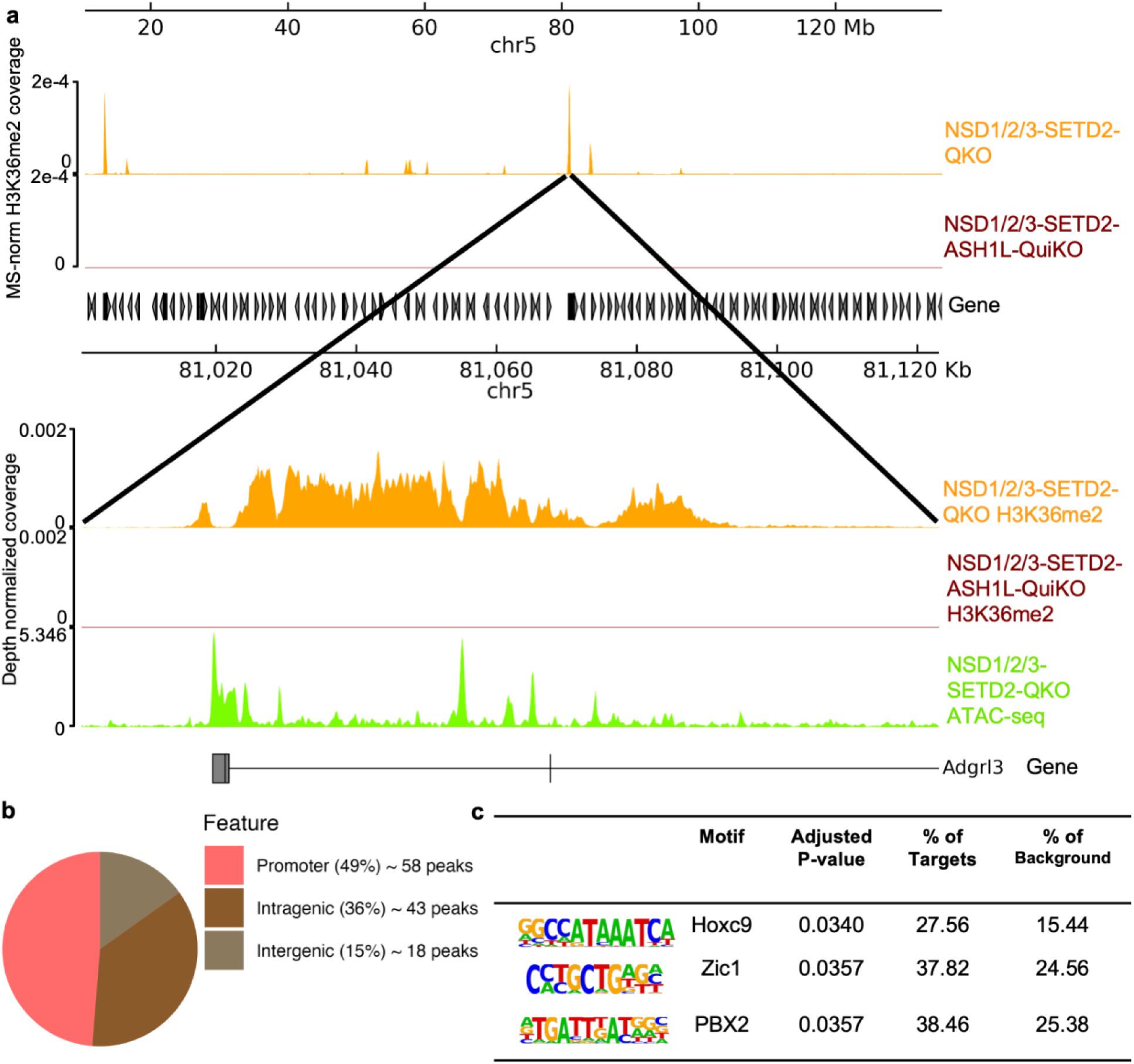
ASH1L-mediated H3K36me2 deposition is targeted to regulatory elements of developmentally-relevant genes. **a** Coverage tracks of MS-normalized H3K36me2 signal for QKO and QuiKO, depicting a region of punctate peaks remaining in QKO cells that are depleted upon subsequent loss of ASH1L. Zooming into one of the peaks shows that they are still quite broad (10-70 kb). **b** Pie chart depicting the distribution of remaining H3K36me2 peaks in QKO cells. Only overlapping peaks from three biological replicates were used in the analysis. **c** Enriched motifs found in open chromatin regions (as marked by ATAC-seq peaks) within H3K36me2 domains in QKO cells, indicating potential recruitment factors for ASH1L.

## Discussion

Over the past decade, a growing body of evidence has demonstrated that H3K36me is essential for the establishment and maintenance of gene regulatory programs (6). Chromosomal abnormalities and loss or gain of function mutations affecting the writers of these marks leads to many different diseases and cancer, underscoring the need to understand how the family of H3K36 methyltransferases regulate their deposition. Our study provides new insights into the distinct genomic preferences of the K36MTs –SETD2, NSD1/2/3, and ASH1L - and helps to consolidate a wealth of previous observations (Fig. 8).

**Fig. 8.**
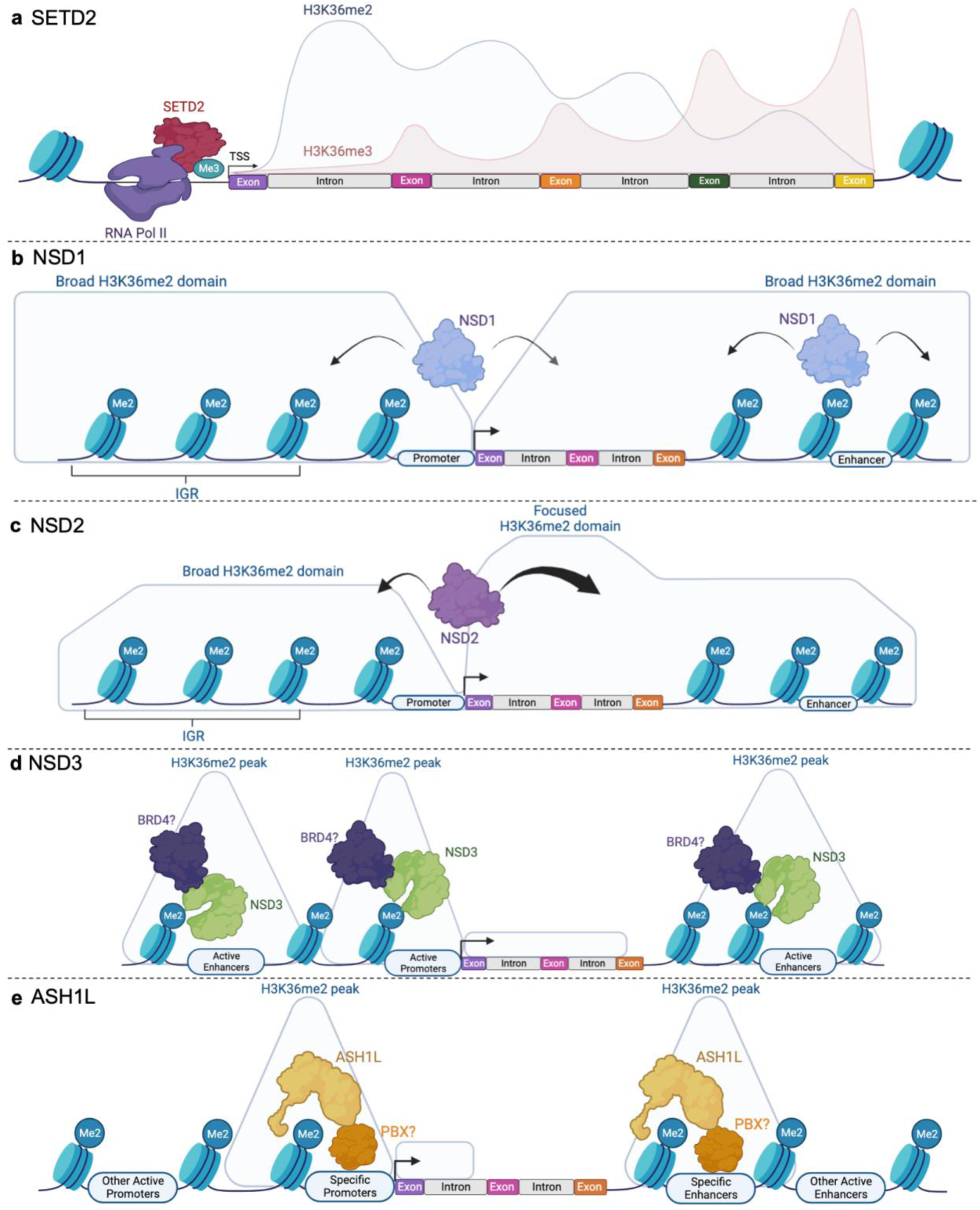
The K36MTs have distinct genomic preferences in the deposition of H3K36me. **a.** SETD2 is coupled to the phosphorylated RNAPII complex (RNA Pol II) and deposits H3K36me3 at exons with higher enrichment in 3’ regions where exon density becomes higher. This takes place at the expense of H3K36me2, which display the opposite 5’ to 3’ and exon/intron trends. **b.** NSD1 is recruited to active CpG islands and then spreads outwards into both gene bodies and intergenic regions, depositing broad H3K36me2 domains. **c.** NSD2 has a specific affinity for depositing H3K36me2 within genes, while also depositing broad intergenic H3K36me2 domains. **d.** NSD3 spreads outwards from CpG-rich regulatory elements and deposits narrower H3K36me2 peaks. **e.** ASH1L primarily deposits H3K36me2 at the promoters and enhancers of specific genes. Created with <u>BioRender.com</u>.

Of the five K36MTs in our study, SETD2 has the most established role in depositing H3K36 methylation. As has been previously reported, it is the dominant H3K36me3- catalyzing enzyme in mammals, but still possesses the ability to deposit H3K36me1/2 (11) (Fig.8). Here, we find that in the absence of SETD2, the levels of H3K36me3 decrease significantly and its genic enrichment disappears. At the same time, the genic distribution of the lower H3K36me2 modification loses its intronic versus exonic enrichment and becomes more correlated with transcription – reflecting its SETD2- independent deposition patterns. This observation highlights the coupling of SETD2 to the phosphorylated RNAPII complex and the transcriptional dependence of its catalytic products. 5’ genic regions generally contain long, rapidly transcribed introns, not allowing SETD2 sufficient time to catalyze the final H3K36me3 step and resulting in predominantly H3K36me1/2 marked nucleosomes. The co-transcriptional process of pre-mRNA splicing slows down the RNAPII complex, thereby allowing more H3K36me3 to be deposited at exons and towards the 3’ ends of genes, as exon density becomes higher. This takes place at the expense of H3K36me1/2, which display the opposite 5’ to 3’ and exon/intron trends. All of the above trends become more pronounced in the absence of NSD1/2, where SETD2 remains the dominant K36MT within gene bodies. Finally, the absence of SETD2 results in an overall increase of H3K36me1/2, reflecting the ability of other enzymes to deposit these two marks and their ineffectiveness in catalyzing the last methylation step.

NSD1 is a demonstrated agent of intergenic H3K36me2 deposition. In HNSCC and mouse embryonic stem cells (mESCs), loss of NSD1 results in a near complete depletion of intergenic H3K36me2 (4,15). Here, in mMSCs, we also find NSD1 to significantly affect intergenic H3K36me2 levels, demonstrating the largest decrease among all individual K36MT KOs. The effects of NSD1 depletion on H3K36me1/3 are much less pronounced. We find that the decrease in H3K36me2 is largest in intergenic regions, however, comparing the single NSD1-KO to NSD1/2-DKO shows that this intergenic depletion is not complete, and that in this cell type, NSD2 and perhaps other enzymes, play a notable role in the deposition of H3K36me2. A recent study demonstrates that NSD1 is recruited to active CGIs, implying that NSD1-associated H3K36me1/2 may then spread outwards into both gene bodies and intergenic regions (Fig. 8) (38). Within gene bodies, it is then upgraded to H3K36me3, suggesting that NSD1 may prime nucleosomes for the deposition of H3K36me3 by providing H3K36me1/2 as the initial substrate.

NSD2 is known to be able to create broad domains of H3K36me2, but this has primarily been shown in the context of oncogenic overexpression (19,39). The catalytic properties of NSD2 under normal physiological conditions have not been well characterized. Here, we find that the individual knockout of NSD2 results in a notable decrease of all H3K36 methylation levels, with the largest fractional decrease occurring for H3K36me3, followed by H3K36me2/1. The depletion of H3K36me2/1 is less pronounced than that observed in NSD1-KO cells, suggesting that similar to other model systems, NSD1 is the dominant H3K36me1/2 methyltransferase operating in euchromatic intergenic regions. However, in mMSCs, it appears that both NSD1 and NSD2 cooperate to produce the intergenic distribution of H3K36me1/2 and the results of individual KOs, as compared to the NSD1/2-DKO is consistent with their additive effect. Interestingly, NSD2-KO appears to result in a slightly greater decrease of H3K36me3, as compared to NSD1-KO. This is consistent with published data showing enrichment of NSD2 within actively transcribed gene bodies (40), leading to the possibility that NSD2 is particularly effective in depositing H3K36me1/2 within genes, possibly – more specifically than NSD1 - acting as a priming substrate for the deposition of H3K36me3 by SETD2 (Fig. 8).

As expected, the NSD1/2-SETD2-TKO cells exhibit very low levels of all three H3K36 methylation marks. However, levels of H3K36me1/2 in particular remain detectable, and a clear enrichment signal is detectable by ChIP-seq. In the case of H3K36me2, there is a remarkable change in the distribution of the signal: while in parental cells H3K36me2 is broadly distributed, and in NSD1/2-DKO cells the signal is mostly restricted to active gene bodies, in TKO cells the signal becomes constrained to far narrower peaks centered on open chromatin and surrounding enhancer and promoter regions. Although the individual KO of NSD3 has little discernible effect on H3K36 methylation, we propose that the other K36MTs can compensate for its absence, and its activity only becomes discernible in the TKO background. As predicted, further deletion of NSD3, creating a QKO cell line results in a near total depletion of all H3K36me levels. Previous work shows that the short isoform of NSD3 is recruited to active regulatory elements by BRD4 (41), and it is likely that the same is true of the full-length isoform of NSD3. Our results are consistent with those observations, suggesting that NSD3, like NSD1, spreads outwards from CpG rich regulatory elements. However, NSD3 is not able to create very broad euchromatic H3K36me2 domains, resulting in the formation of much narrower peaks (Fig. 8).

Finally, we find ASH1L to be the least prolific, and most specific K36MT. Similar to NSD3, the individual knockout of ASH1L has no detectable effect on H3K36me levels and distribution, implying that in this cell type, other K36MTs can compensate for its loss. By comparing the QKO to the QuiKO cells, we show that ASH1L is only responsible for depositing H3K36me2 at the promoters and enhancers of approximately 100 developmentally related genes, and its ability to spread the modification is even lower than that of NSD3 (Fig. 8). It remains to be seen what mechanisms govern the recruitment of ASH1L to those specific regions. TFBS analysis suggests that specific transcription factors, particularly from the PBX family may be involved.

Our results illuminate some of the intricate dependencies governing the K36MTs and the deposition of H3K36 methylation. The overall outcomes depend on several key variables: genomic recruitment and the distribution of K36MTs; their ability to spread and read existing modifications through their PWWP domains; their catalytic activities; and finally, expression levels in various cellular contexts. Many of these parameters are being continually refined. In this study, we find a hierarchy of K36MT activities pertaining to the deposition of H3K36me1/2, with NSD1>NSD2>NSD3>ASH1L. We find that in contrast to mESCs, the function of NSD1 is less prominent and NSD2 contributes significantly to the normal deposition of H3K36me2. SETD2 retains its position as the dominant H3K36me3 methyltransferase. Furthermore, trace levels of H3K36me1 remain in the QuiKO cells, and the distribution patterns largely mirror those observed in parental cells. Other protein families outside of the scope of this study have also been implicated in the deposition of H3K36me, such as the SMYD family (42,43), and this residual H3K36me1 indicates that another enzyme may be responsible for its deposition, albeit at extremely low levels.

H3K36 methylation also has profound impacts on the deposition of other histone marks and DNAme. In the context of active transcription, H3K36me2/3 coexist with marks such as H3K4me1/3, H3K27ac, and DNAme either within genes and/or at CREs. Conversely, H3K36me2/3 tend not to coexist with marks that define constitutive or facultative heterochromatin, such as H3K9me3 and H3K27me3, respectively. In the presence of mutations affecting the deposition of H3K36 methylation, such as H3K36M, the downstream consequences affect the establishment and maintenance of all these marks, and in turn result in aberrant gene expression (4,14,15,31). Therefore, through better understanding the regulation of H3K36 methylation, our work helps pave the way for future studies to further define these and other relationships that exist between epigenetic marks.

## Conclusions

Here, we identify both unique and overlapping roles by the family of K36MTs in their regulation of H3K36me. Our findings demonstrate that within transcribed genes, SETD2 deposits both H3K36me2/3, while the other K36MTs are capable of depositing H3K36me1/2 independently of SETD2 activity. In most cell types, NSD1 is responsible for most propagation of intergenic H3K36me2, however, in mMSCs, NSD2 also contributes to its deposition in these regions. We also identified that the catalytic products of NSD3 are primarily focused on active promoters and enhancers, and that the activity of ASH1L is even more restricted to specific developmental genes and their regulatory elements. Given their implications in both development and disease, our study helps to illuminate how the family of K36MTs collectively regulate the deposition of H3K36me. Despite the overlap in the regions where each K36MT exerts its catalytic functions, they each have unique properties in the propagation of these marks.

## Materials and methods

### Cell culture, CRISPR-Cas9 gene editing and generation of stable cell lines

C3H10T1/2 mouse mesenchymal stem cells (ATCC) were cultured in DMEM with 10% fetal bovine serum (FBS,) and supplemented with 1% GlutaMax. *Drosophila* S2 cells were cultured in Schneider’s *Drosophila* medium (ThermoFisher) containing 10% heat-inactivated FBS. All cell lines tested negative for mycoplasma contamination. To generate knockout cell lines, 3.5 x 10^5^ C3H10T1/2 cells were nucleofected with ribonucleoprotein (RNP)-mediated CRISPR-Cas9 using the Alt-R CRISPR-Cas9 System (IDT) and Lonza Amaxa® SE Cell Line 4D Nucleofector kit (V4XC-1032, program CA-133). Synthetic crRNA guides were designed (Supplementary Table 1) and combined with Alt-R® CRISPR-Cas9 tracrRNA, ATTO 550 (IDT) and coupled to Alt-R® S.p. Cas9 Nuclease V3 following the IDT protocol “Delivery of ribonucleoprotein complexes using the Lonza® Nucleofector System” prior to nucleofection. The transfected cells were incubated for 48h. Single ATTO 550 positive cells were then sorted into 96-well plates. Individual clones were then expanded and validated by MiSeq sequencing using specific primers for target loci (Supplementary Table 1). Three biological clones were chosen and expanded for each knockout cell line, and used in all subsequent assays. C3H10T1/2 cells overexpressing the H3K36M mutation were provided by the lab of Dr. Chao Lu (Stanford University), and C3H10T1/2 NSD1/2-DKO and C3H10T1/2 NSD1/2-SETD2-TKO cells were provided by the lab of Dr. C. David Allis (Rockefeller University).

### Crosslinking and ChIP-seq

Approximately 20 million mMSC cells per cell line were used. Crosslinking was performed in 150 mm cell culture plates using 1% formaldehyde (Sigma) at room temperature with gentle rocking for 10 minutes. The crosslinking reaction was quenched using 1.25 M Glycine with gentle rocking at room temperature for 5 minutes. Fixed cell preparations were washed with ice-cold PBS, scraped, and then washed twice more with ice-cold PBS. Crosslinked pellets were resuspended in 500 μl cell lysis buffer (5 mM pH 8.5 PIPES, 85 mM KCl, 1% (v/v) IGEPAL CA-630, 50 mM NaF, 1 mM PMSF, 1 mM phenylarsine oxide, 5 mM sodium orthovanadate, EDTA-free protease inhibitor tablet) and incubated for 30 minutes on ice. Samples were centrifuged and pellets were resuspended in 500 μl of nuclei lysis buffer (50 mM pH 8.0 Tris-HCl, 10 mM EDTA, 1% (w/v) SDS, 50 mM NaF, 1 mM PMSF, 1 mM phenylarsine oxide, 5 mM sodium orthovanadate, EDTA-free protease inhibitor tablet) and incubated for 30 minutes on ice. Sonication of lysed nuclei was performed using the BioRuptor UCS-300 at maximum intensity for 75-90 cycles (10s on, 20s breaks). Sonication efficiency to achieve fragments between 150 - 500 bp was evaluated using gel electrophoresis on a reversed-crosslinked and purified aliquot from each sample. After sonication, chromatin was diluted to reduce SDS levels to 0.1% and concentrated using Nanosep 10K OMEGA (Pall) columns. Prior to the chromatin immunoprecipitation (ChIP) reaction, 2% sonicated *Drosophila* S2 cell chromatin was spiked into each sample for quantification of total levels of histone marks. The ChIP reactions were performed using the Diagenode SX-8G IP-Star Compact and Diagenode automated iDeal ChIP-seq Kit for Histones. Dynabeads Protein A (Invitrogen) were washed, then incubated with specific antibodies (Supplementary Table 2), 1.5 million cells of sonicated cell lysate, and protease inhibitors for 10 hrs, followed by a 20 min wash cycle using the provided wash buffers (Diagenode Immunoprecipitation Buffers, iDeal ChIP-seq Kit for Histones). Reverse cross-linking was then performed using 5M NaCl at 65℃ for 4 hrs. ChIP samples were then treated with 2 μl RNase Cocktail at 65℃ for 30 min followed by 2 μl Proteinase K at 65℃ for 30 min. Samples were then purified with QIAGEN MinElute PCR purification kit (QIAGEN) as per the manufacturer’s protocol. In parallel, input samples (chromatin from about 50,000 cells) were reverse crosslinked and DNA was isolated following the same protocol. Library preparation was performed using the Kapa Hyper Prep library preparation reagents following the manufacturer’s protocol (Kapa Hyper Prep Kit, Roche 07962363001). ChIP libraries were sequenced using the Illumina HiSeq 4000 at 50 bp single reads or NovaSeq 6000 at 100 bp single reads.

### Visualization

Unless otherwise stated, figures were created using ggplot2 (44) v3.3.0. Coverage/alignment tracks were visualized using pyGenomeTracks (45) v3.2.1.

### ChIP-seq processing and analysis

Publicly-available NSD1/2-SETD2-TKO H3K27ac ChIP-seq data was used in this study (37).

Reads were processed as described (4), where they were aligned using BWA (46) to a combined reference of mm10 and dm6 and afterwards filtered using a cut-off of MAPQ < 3 using Samtools (47). Samclip v.0.2 (samtools view -h in.bam | samclip --ref ref.fa | samtools sort > out.bam) was used to filter contaminated sequences for H3K36me2 NSD1/2/3-SETD2-QKO and NSD1/2/3-SETD2-ASH1L-QuiKO replicates prior to alignment (48).

Raw tag counts were binned into windows using bedtools (49) v2.29.0 as previously described (15) in different sized bins (1kb and 10kb). deepTools (50) v3.3.1 was used to normalize ChIP signals either by dividing the total alignments (in millions) (bamCoverage -b $BAM -o $OUTPUT.bigWig --normalizeUsing CPM --centerReads -e 200) or additionally taking the log2 ratio of ChIP signals by those of the input (bamCompare -b1 $BAM_ChIP -b $BAM_Input -o OUTPUT.bigWig -e 200 –centerReads -bs 200 --blackListFileName $BL_bed --normalizeUsing CPM --scaleFactorsMethods None). PCA and hierarchical clustering based on the Pearson’s correlation matrix were generated using the log2 ratio of ChIP signals over input values from 10kb bins. To generate coverage tracks, following CPM normalization as indicated above, replicates were merged in a stepwise fashion using bigwigCompare from deepTools with parameters ‘-b1 rep1 -b2 rep2 $outdir --operation mean -bs 200 -o $merged.step1.cpm.bw’ and ‘-b1 merged.step1.cpm.bw -b2 rep3 $outdir --operation mean -bs 200 -o $merged.final.cpm.bw’. Additionally, quantitative normalization entailed multiplying the original signal (either in CPM or log2 ratio over input) by the genome-wide modification percentage values obtained from mass spectrometry (these values were averaged if normalizing merged bigwig tracks), as previously described (15). The ENCODE blacklist (51) was used.

Gene body distribution plots were generated using computeMatrix using genes with at least 50000 bp width and an exon count of at least 6. Furthermore, to avoid low gene counts, only the top 50th percentile (mean counts > 0.012 after normalization using DESeq2 and adjusted for gene length). Only the first and last three exon/intron pairs are shown in the plots. Annotated promoters were defined as 1000 bp upstream of the transcription start sites. The depth-normalized mean signal found in each annotation was calculated using computeMatrix with parameters ‘scale-regions -R $feature.bed -S $sample.bw -bl $BL_bed -b 0 -a 0 -m 1000 -bs 5 --samplesLabel $sample_name’. Correlation plots depicting the relationship between H3K36me2 and gene expression quantiles were created by plotting the average depth-normalized H3K36me signal within gene bodies, taking into account the exclusion criteria mentioned above. This was plotted against the average gene expression signal within each quantile.

Enrichment matrices for aggregate plots and heatmaps were generated using computeMatrix from deepTools for intergenic regions (scale-regions --regionBodyLength 20000 --beforeRegionStartlength 20000 --afterRegionStartLength 20000 --binSize 1000), for promoters (reference-point -R $Promoters -bl $BL_bed --referencePoint center -- binSize 50 -a 3000 -b 3000 --missingDataAsZero --skipZeros) and for enhancers (reference-point -R $Enhancers -bl $BL_bed --referencePoint center --binSize 50 -a 3000 -b 3000 --missingDataAsZero --skipZeros). Genic regions were taken as the union of any intervals having the “gene” annotation in Ensembl and intergenic regions thus defined as the complement of genic ones. The ratio of intergenic enrichment over neighboring genes was calculated as previously described (15). The exon-to-intron ratio was determined by utilizing the deepTools plotEnrichment tool to quantify H3K36me levels in annotated exons and introns sourced from Ensembl. Subsequently, the counts were normalized based on sample library depth and total length of annotations. Active enhancers were identified as those containing parental/wildtype ATAC-seq peaks, while inactive enhancers lacked such peaks. To ensure equal numbers of active and inactive enhancers, random sampling without replacement was performed, resulting in a sample size of n = 1758 for active enhancers and n = 1758 for inactive enhancers. computeMatrix using the parameters ‘reference-point -R $inactive_enhancers $active_enhancers -bl $BL_bed --referencePoint center --binSize 50 -a 3000 -b 3000 --missingDataAsZero -- skipZeros --samplesLabel $samples_label -o $mat.gz’. The same parameters for computeMatrix were used to generate heatmaps for inactive and active promoters.

“Peakiness” scores were computed as the average read-depth normalized coverage of the top 1% most covered 1-kb windows across the genome, excluding those overlapping blacklisted regions, as previously described (4).

10 kb bins with H3K36me2 signal in NSD1/2-SETD2-TKO not found in NSD1/2/3- SETD2-QKO were used for overlap enrichment analysis. Overlap enrichment was determined with all the bins as the background set as implemented in LOLA (52) v1.16.0 for Ensembl (35) 97 annotations (genes and regulatory build (53)).

SPAN (54) v1.1, a semi-supervised peak-caller, was used to call peaks for NSD1/2/3-SETD2-QKO samples (java -Xmx8G -jar span.jar analyze -t $ChIP.bam -c $Control.bam --cs $mm10.chrom.sizes -p $Results.peak). ChIPseeker was applied (with options: genomicAnnotationPriority = c(“Intergenic”,“Promoter”, “Exon”,“Intron”), tssRegion = c(−1000, 1000)) to obtain the genomic feature distribution for the intersect of peaks from NSD1/2/3-SETD2-QKO replicates (55).

Motifs were obtained using HOMER (56) v4.11 for ATAC-seq peaks within H3K36me2 peaks in the intersection of peaks from NSD1/2/3-SETD2-QKO replicates.

To generate genome-wide correlation plots between genes and H3K36me2 signal, we utilized featureCounts (57) (version 1.5.3) to count H3K36me2 reads within exons, which were then aggregated for all exons belonging to their corresponding genes. Subsequently, the raw counts underwent normalization using DESeq2’s median of ratios method (58), scaled according to the total length of exons for each gene and log transformed after removing any normalized read counts equal to zero.

ChIP-Rx was processed as previously described (31). ChIP-Rx was calculated for each sample by dividing the ratio of the total ChIP-seq reads mapping to the mm10 over the total ChIP-seq reads mapping to dm6 over the ratio of the total input reads mapping to mm10 over the total input reads mapping to dm6 (see below). See Supplementary Table 3 for calculated ChIP-Rx values.

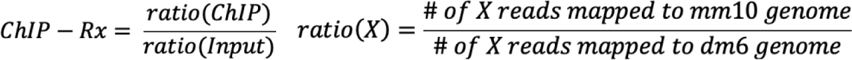

### ATAC-seq

ATAC-Seq library preparation was performed according to the Omni-ATAC protocol (59). 50,000 C3H10T1/2 cultured cells were resuspended in 1 ml of cold ATAC-seq resuspension buffer (RSB; 10 mM Tris-HCl pH 7.4, 10 mM NaCl, and 3 mM MgCl2 in water). Cells were centrifuged at 500 rcf for 5 min in a pre-chilled (4°C) fixed-angle centrifuge. After centrifugation, supernatant was aspirated and cell pellets were then resuspended in 50 μl of ATAC-seq RSB containing 0.1% IGEPAL, 0.1% Tween-20, and 0.01% digitonin by pipetting up and down three times. This cell lysis reaction was incubated on ice for 3 min. After lysis, 1 ml of ATAC-seq RSB containing 0.1% Tween-20 (without IGEPAL and digitonin) was added, and the tubes were inverted to mix. Nuclei were then centrifuged for 10 min at 500 rcf in a pre-chilled (4°C) fixed-angle centrifuge. Supernatant was removed and nuclei were resuspended in 50 uL transposition mix (2x TD Buffer, 100 nM final transposase, 16.5 uL PBS, 0.5 uL 1% digitonin, 0.5 uL 10% Tween-20, 5 uL H2O) (Illumina Tagment DNA Enzyme and Buffer Small Kit, 20034197). Transposition reactions were incubated at 37 °C for 30 min in a thermomixer with shaking at 1000 rpm. Reactions were cleaned up with DNA Clean and Concentrator 5 columns (Zymo Research). Illumina Nextera DNA Unique Dual Indexes Set C (Illumina, 20027215) were added and amplified (12 cycles) using NEBNext 2x MasterMix. Sequencing of the ATAC-Seq libraries was performed on the Illumina NovaSeq 6000 system using 100-bp paired-end sequencing.

### ATAC-seq processing and analysis

Publicly available parental/wildtype ATAC-Seq data from Rajagopalan et al. (2021) (37) was used to distinguish between inactive and active enhancers. Raw reads were trimmed and filtered for quality using Trimmomatic v0.39 (60). Trimmed reads were mapped to the mm10 genome assembly using Bowtie2 v2.5.1 (61), and non-uniquely mapping reads were removed. Afterwards, the reads were adjusted by shifting all positive-strand reads 4 bp downstream and all negative-strand reads 5 bp upstream to the center of the reads on the transposase binding event. Peak calling was performed on each replicate using MACS2 v2.2.6 (62) with ‘-extsize 200 -shift -100 -nomodel’ parameters. To find a set of reproducible peaks across replicates, we calculated the irreproducible discovery rate (IDR) (63) and excluded peaks with an IDR greater than 0.05 across every pair of replicates. Subsequently, the ENCODE blacklist (51) was used to filter the peaks. Coverage tracks were generated using bamCoverage with parameters ‘-b $BAM -o $OUTPUT.bigWig --normalizeUsing CPM --centerReads -e 200 --minMappingQuality 5 - bs 200’’.

### RNA-seq processing and analysis

Publicly available RNA-seq data in mMSC was used for this study, specifically three replicates of parental C3H10T1/2 mouse mesenchymal stem cells (37). Raw reads were aligned to mm10 genome build using STAR version 2.5.3a (64). Afterwards, featureCounts (57) (version 1.5.3) was used to count exonic reads from the GTF annotation (GENCODE version from UCSC). Expression was normalized using DESeq2’s median of ratios (58), and divided by the aggregate length of exons per gene. GENCODE VM25 gene annotation was used to filter for transcripts that are at least 50000 bp and have at least 6 exons. Genes consisting of these transcripts were categorized into five groups based on their expression levels, which was calculated by taking the average of parental replicates. After filtering for genes below the 50th percentile (normalized counts greater than 0.01), the remaining genes were divided into four quantiles and hence four groups of genes based on expression level. A fifth group of genes were designated that had zero expression. The transcripts for each of these genes were then further subdivided into 10 kb upstream of the promoter, promoter region, the first three exon/intron pairs, the last three exon/introns pairs, and lastly, 10 kb downstream of the last exon. Enrichment matrices were then generated for each feature and for each gene expression quantile using deepTools’ computeMatrix (computeMatrix scale-regions -R $Feature.bed -S $Sample.bigWig -bl $BL.bed -b 0 -a 0 -m 1000 -bs 5 -o $output.mat.gz).

For obtaining “Active promoters” and “Inactive promoters”, the remaining genes after filtering for zero reads were divided into three quantiles. The four groups were randomly downsampled to the group with the lowest number of genes. Finally, promoters were obtained using GenomicRanges::promoters() v3.17 (65) for these four groups of genes. Here, the “Active promoters” are referred to as an aggregate of the top two groups with the highest gene expression and hence most active promoters, “Inactive promoters” are the remaining two groups with low gene expression and least active promoters. To ensure equal numbers of active and inactive promoters, random sampling without replacement was performed, resulting in a sample size of n = 18,290 for active promoters and n = 18,290 for inactive promoters.

### Histone acid extraction, histone derivatization, and analysis of post-translational modifications by nano-LC-MS

4 million cells from each clonal mMSC line were collected and frozen at -80℃. Thawed pellets were lysed in nuclear isolation buffer (15 mM Tris pH 7.5, 60 mM KCl, 15 mM NaCl, 5 mM MgCl2, 1 mM CaCl2, 250 mM sucrose, 10 mM sodium butyrate, 0.1% (v/v) beta-mercaptoethanol, commercial phosphatase and protease inhibitor cocktail tablets) containing 0.3% NP-40 alternative on ice for 5 min. Nuclei were subsequently washed twice in the same buffer without NP-40, and pellets were resuspended using gentle vortexing in chilled 0.4 N H2SO4, followed by a 3 hr incubation while rotating at 4℃. After centrifugation, supernatants were collected and proteins were precipitated in 20% TCA overnight at 4℃, washed with 0.1% HCl (v/v) acetone once, followed by two washes with acetone alone. Histones were resuspended in deionized water. Acid-extracted histones (20 μg) were resuspended in 50 mM ammonium bicarbonate (pH 8.0), derivatized using propionic anhydride and digested with trypsin as previously described (15). After the second round of propionylation, the resulting histone peptides were desalted using C18 Stage Tips, dried using a centrifugal evaporator and reconstituted using 0.1% formic acid in preparation for LC-MS analysis. Nanoflow liquid chromatography was performed using a Thermo Fisher Scientific, Vanquish Neo UHPLC equipped with an Easy-Spray™ PepMap™ Neo nano-column (2 µm, C18, 75 µm X 150 mm). Buffer A was 0.1% formic acid and Buffer B was 0.1% formic acid in 80% acetonitrile. Peptides were resolved using at room temperature with a mobile phase consisting of a linear gradient from 1 to 45% solvent B (0.1% formic acid in 100% acetonitrile) in solvent A (0.1% formic acid in water) over 85 mins and then 45 to 98% solvent B over 5 mins at a flow rate of 300 nL/min. The HPLC was coupled online to an Orbitrap Exploris 240 (Thermo Scientific) mass spectrometer operating in the positive mode using a Nanospray Flex Ion Source (Thermo Fisher Scientific) at 1.9 kV. A full MS scan was acquired in the Orbitrap mass analyzer across 350–1050 m/z at a resolution of 120,000 in positive profile mode with an auto maximum injection time and an AGC target of 300%. Parallel reaction monitoring experiments were followed for monitoring the targeted peptides based on the inclusion list. Targeted ions were fragmented using HCD fragmentation. These scans typically used an NCE of 30, an AGC target standard, and an auto maximum injection time. Raw files were analyzed using EpiProfile 2.0 (66) and Skyline (67). The area for each modification state of a peptide was normalized against the total signal for that peptide to give the relative abundance of the histone modification. See Supplementary Tables 4 and 5 for raw m/z data per modification and calculated MS values, respectively.

### Western blotting

For each C3H10T1/2 Western blot sample, 1 million cells were collected and counted using the automated Countess II cell counter (ThermoFisher). Each cell pellet was washed with PBS. For histone marks, each pellet was resuspended in 100 μl of 1X Laemmli lysis buffer, 1:100 proteinase inhibitor cocktail, and 0.1 mM PMSF (Laemmli 6X stock contains 0.35M Tris-HCl pH 6.8, 30% glycerol, 10% SDS, 20% beta-mercaptoethanol, 0.04% bromophenol blue, in water). Samples were then sonicated using the Bioruptor UCD-300 for 10 cycles at max intensity (30s on, 20s breaks), and ran on stain-free TGX 4%-15% gradient pre-cast gels (Bio-Rad, 4568084). Semi-dry electrotransfer onto a PVDF membrane was completed using the trans-blot Turbo Transfer system (Bio-Rad, Trans-blot® Turbo RTA Mini LF PVDF Transfer Kit, 1704274). PVDF membranes were blocked with 5% BSA in TBSt, and probed overnight with the respective primary antibody (Supplementary Table 2). Following washes with TBSt, membranes were incubated for 1h with anti-rabbit horseradish peroxidase-conjugated secondary antibody in 2% BSA-TBSt. Imaging was completed using the ECL Clarity or Clarity Max solutions (Bio-Rad, 1705060, 1705062). Relative intensities quantified using the Bio-Rad Image Lab™ Software (RRID:SCR_014210).

### Antibodies

Antibodies:
|  |  |  |
| --- | --- | --- |
| Rabbit monoclonal anti-H3K36me1 | Cell Signaling Technology | D9J1D, Cat # 14111 |
| Rabbit monoclonal anti-H3K36me2 | Cell Signaling Technology | C75H12, Cat # 2901 |
| Rabbit polyclonal anti-H3K36me3 | Active Motif | RRID:AB_2615073, Cat # 61101 |
| Rabbit polyclonal anti-H3K27ac | Diagenode | RRID:AB_2637079, Cat # C15410196 |

## Declarations

### Ethics approval and consent to participate

Not applicable.

### Consent for publication

Not applicable.

### Availability of data and materials

The publicly-available NSD1/2-SETD2-TKO H3K27ac ChIP-seq data, Parental ATAC- seq and Parental RNA-seq data were obtained from the National Center for Biotechnology Information Gene Expression Omnibus with accession number GSE160266 (37). Newly generated mMSC sequencing data (ChIP-Seq and ATAC-Seq) are available in the National Center for Biotechnology Information Gene Expression Omnibus with accession number GSE243566. Link to access data: https://www.ncbi.nlm.nih.gov/geo/query/acc.cgi?acc=GSE243566.

## Authors’ contributions

G.S. and R.P. contributed equally to this work. Study design, G.S., R.P., and J.M. Writing (original draft), G.S., R.P., and J.M. Laboratory experiments, G.S. Bioinformatic analyses, R.P. and B.H. Data processing, R.P. and E.B. Mass spectrometry and subsequent analysis, F.V. and J.G. Supervision, J.M., B.G., and C.L. Funding acquisition, J.M., B.G., and C.L. All author(s) read and approved the final manuscript.

## Acknowledgements

We would like to thank all members of the J.M., B.G., and C.L. groups for their input and feedback. We thank the McGill University Genome Centre for their expertise in sample quality verification and sequencing services. We thank the group of C.L. for generation of C3H10T1/2 cells overexpressing the H3K36M mutation and Dr. Daniel Weinberg for earlier work that generated many of the resources that became the core of this research. We owe an unmeasurable debt of gratitude to Dr. David Allis who, although no longer with us, continues to inspire our investigations of chromatin biology.

## Supplementary figures

**Fig. S1.**
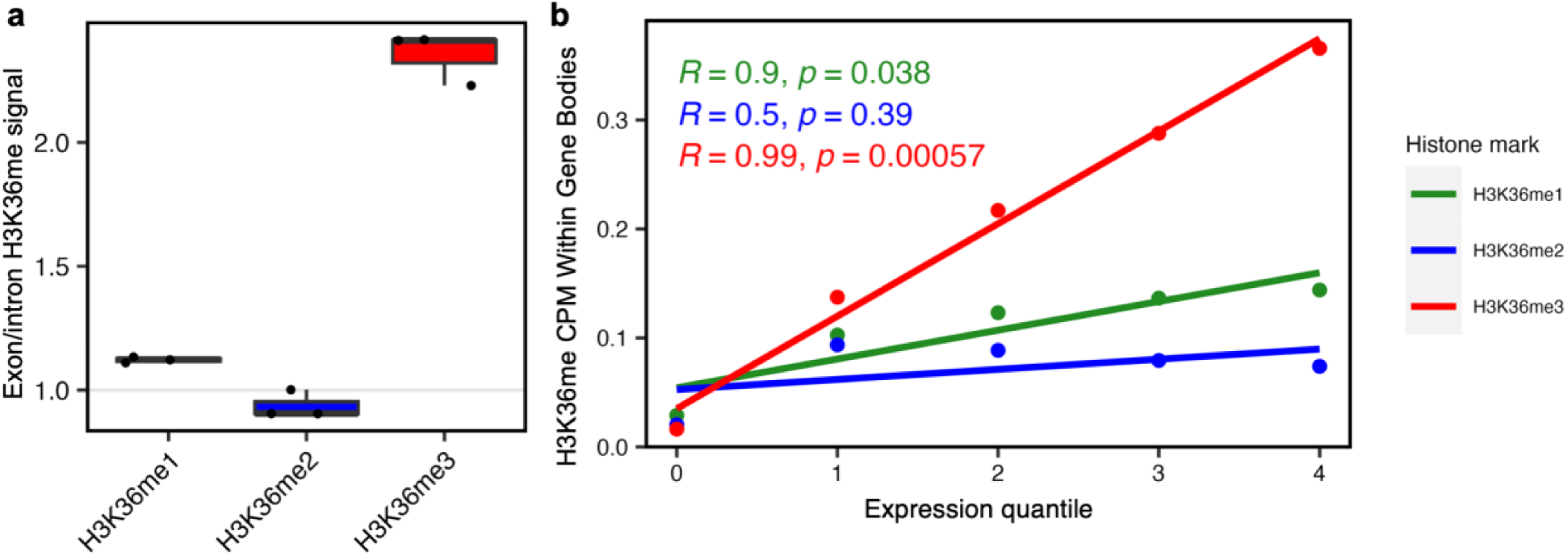
Genome-wide characterization of H3K36me indicates distinct patterns within gene bodies and dependence on transcription. **a**. Boxplots of H3K36me1/2/3 enrichment at exons versus introns, illustrating higher H3K36me1 and H3K36me3 signals at exons whereas the inverse is found for H3K36me2. **b.** Correlation plots illustrating the relationship between H3K36me1/2/3 signal and gene expression quantiles. Reported values are Pearson’s correlation coefficient (*R)*.

**Fig. S2.**
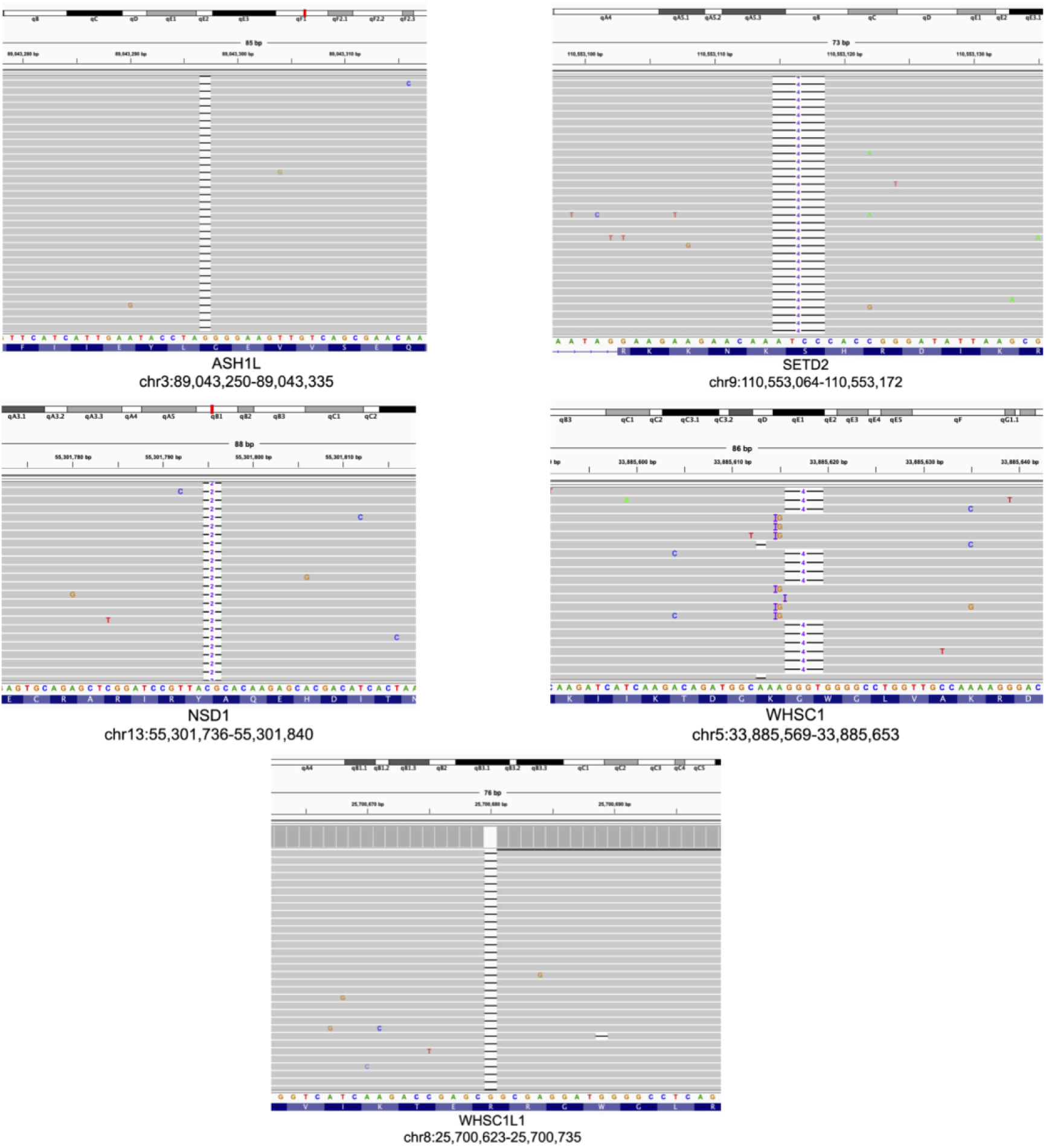
Representative genome-browser coverage tracks validating CRISPR gene editing as assessed by MiSeq. gRNAs were designed to target the catalytic SET domain of each H3K36 methyltransferase. Gene names and genomic coordinates of edits are indicated at the bottom of each set of coverage tracks. WHSC1 and WHSC1L1 are the mouse orthologs of human NSD2 and NSD3, respectively. Similar edits were achieved in multi-knockout conditions (i.e. NSD1/2/3-SETD2-QKO and NSD1/2/3-SETD2-ASH1L-QuiKO). Three biological clones were obtained for each condition.

**Fig. S3.**
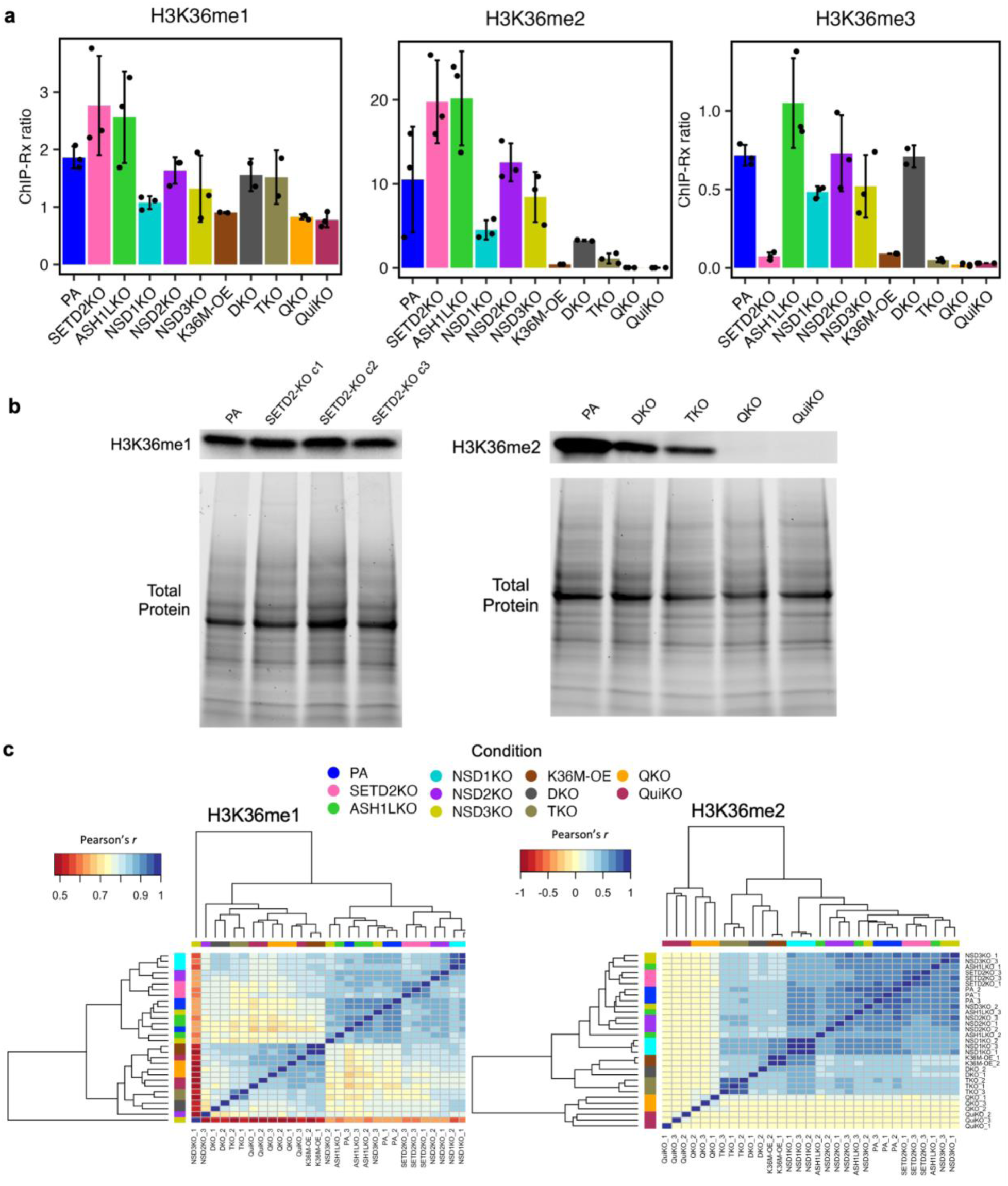
Various K36MT deletions lead to distinct genome-wide alterations to H3K36me. **a**. Global abundance of H3K36me1/2/3 as quantified using ChIP-seq with *Drosophila* spike-in reference chromatin (ChIP-Rx) (as described in methods). Error bars represent mean ± standard deviation. **b.** Western blots for H3K36me1 (left) and H3K36me2 (middle). Total protein loaded and used for quantification shown below each blot. Western blot comparing parental and SETD2-KO H3K36me1 levels (right). Quantification performed as described in methods. **c.** Pearson’s correlation matrix-based hierarchical clustering of H3K36me1/2, illustrating the clustering of single knockouts with PA whereas H3K36M-OE clusters with the multiple knockouts.

**Fig. S4.**
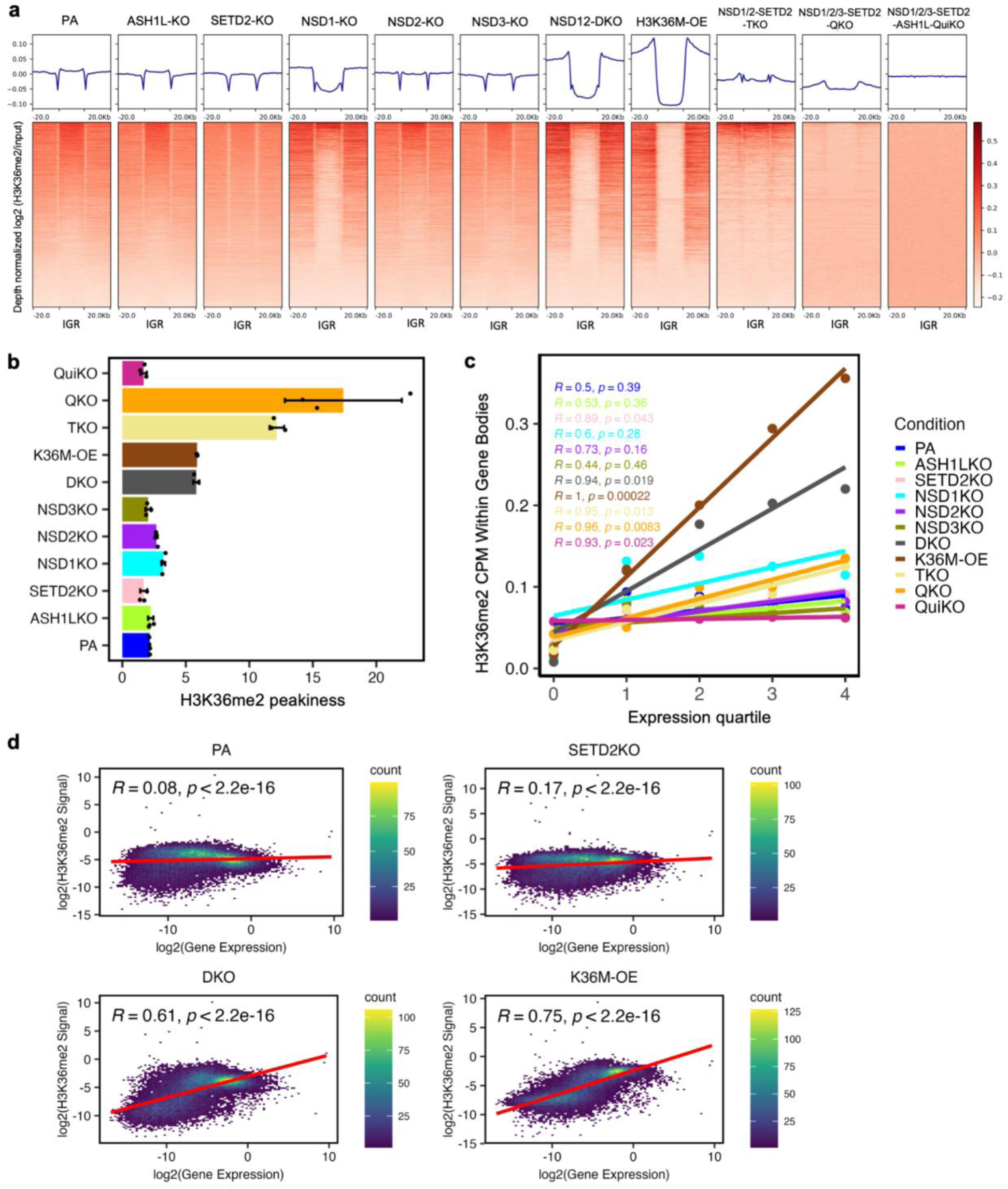
Distinct H3K36me2 distribution patterns are found following knockout of K36MTs. **a.** Heatmaps showing H3K36me2 (input- and depth-normalized) enrichment patterns ± 20 kb flanking IGRs, summarizing the H3K36me2 trends at IGR in parental and knockout cells. **b.** H3K36me2 “peakiness score”, representing the average ChIP signal in the top 1% of 1-kb bins. More punctate distributions of H3K36me2 are found following multiple deletions whereas individual knockouts have less effect on peakiness score. Individual data points represent replicates for each condition. **c.** Correlation plots illustrating the relationship between H3K36me2 signal and gene expression quantiles. Reported values are Pearson’s correlation coefficient (*R).* With the exception of SETD2-KO, single knockouts have no effect on the correlation between H3K36me2 and gene expression. **d.** Genome-wide correlation analysis between exonic H3K36me2 and gene expression, illustrating stronger correlations following SETD2-KO, NSD1/2-DKO and H3K36M-OE compared to parental. Only genes with H3K36me2 and gene expression counts greater than zero were included in the analysis. Both the H3K36me2 and gene expression exonic read counts (aggregated per gene) were normalized using DESeq2, scaled based on the cumulative length of all exons within each gene, and then log transformed.

**Fig. S5.**
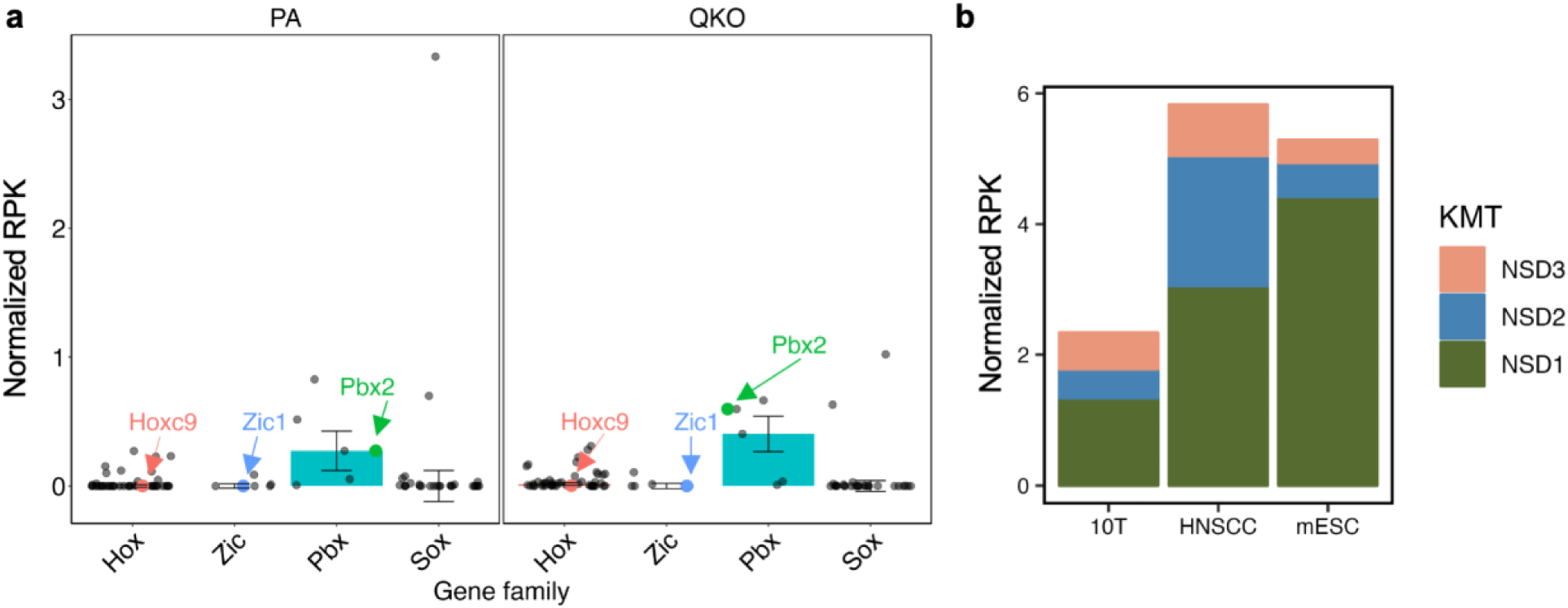
NSD1/2/3 are expressed at different relative levels across three different cell lines. **a-b** Bar graphs showing DESeq2 counts that have been normalized, with each gene scaled according to its specific gene length. **a.** Only the Pbx gene family, including Pbx2, is expressed in 10T-mMSCs, as shown in both parental and NSD1/2/3-SETD2-QKO cell lines. Error bars represent median ± standard error. **b.** Relative expression of NSD1/2/3 in mouse mesenchymal stem cells (mMSCs), mouse embryonic stem cells (mESCs), and human head and neck squamous cell carcinoma (HNSCC) cells.

